# Skull evolution and lineage diversification in endemic Malagasy carnivorans

**DOI:** 10.1101/2024.03.25.586658

**Authors:** Chris J. Law, Tate J. Linden, John J. Flynn

**Affiliations:** Department of Integrative Biology, University of Texas; Burke Museum and Department of Biology, University of Washington; Division of Paleontology, American Museum of Natural History

**Keywords:** **Key terms**: Adaptive radiation, Carnivora, craniomandibular, Madagascar, phenotypic evolution, phylogenetic comparative methods

## Abstract

Madagascar is one of the world’s foremost biodiversity hotspots with more than 90% of its species endemic to the island. Malagasy carnivorans are one of only four extant terrestrial mammalian clades endemic to Madagascar. Although there are only eight extant species, these carnivorans exhibit remarkable phenotypic and ecological diversity that is often hypothesized to have diversified through an adaptive radiation. Here, we investigated the evolution of skull diversity in Malagasy carnivorans and tested if they exhibited characteristics of convergence and an adaptive radiation. We found that their skull disparity exceeds that of any other feliform family, as their skulls vary widely and strikingly capture a large amount of the morphological variation found across all feliforms. We also found evidence of shared adaptive zones in cranial shape between euplerid subclades and felids, herpestids, and viverrids. Lastly, contrary to predictions of adaptive radiation, we found that Malagasy carnivorans do not exhibit rapid lineage diversification and only marginally faster rates of mandibular shape evolution, and to a lesser extent cranial shape evolution, compared to other feliforms. These results reveal that exceptional diversification rates are not necessary to generate the striking phenotypic diversity that evolved in carnivorans after their dispersal to and isolation on Madagascar.

## Introduction

Many islands are considered as biodiversity hotspots [1] and contain species that are not found anywhere else in the world [2–4]. High rates of endemism could be spurred by adaptive radiations [5]. Islands in particular, which are often geographically isolated, provide novel ecological opportunities for founding species to rapidly diversify to a variety of species and phenotypes to fill previously unoccupied ecological niches [6–8]. Unsurprisingly, the best-studied adaptive radiations are found among insular island groups such as Galápagos finches, Hawaiian honeycreepers, and Caribbean anoles [9–11]. One of the world’s foremost biodiversity hotspots is Madagascar, with more than 90% of its species endemic to the island [12]. As Madagascar has been geographically isolated for over 88 million years, most endemic Malagasy vertebrates are a result of several independent successful dispersal events from Africa or Asia [13]. Upon arrival on Madagascar, adaptive radiations are hypothesized to have facilitated the diversification of many Malagasy clades, such as beetles [14,15], frogs [16,17], vangas [18], and lemurs [19–21]. In this study, we investigate the diversity of Malagasy euplerid carnivorans, one of four extant terrestrial mammalian clades (i.e., Lemuroidea, Eupleridae, Tenrecidae and Nesomyinae) endemic to Madagascar and hypothesized to have diversified through an adaptive radiation [22].

Although there are only eight extant species, Malagasy carnivorans exhibit remarkable phenotypic and ecological diversity [22–25], so much so that no anatomical character can be used to define them. As a result, Malagasy carnivorans have traditionally been thought to belong to or originate from multiple feliform families (i.e., Herpestidae, Viverridae, and Felidae) and thus presumably dispersed to Madagascar through multiple independent dispersal events [26–29]. Molecular data has since revealed that all Malagasy carnivorans originated from a single ancestor, thus forming a monophyletic clade and most likely exhibited a single dispersal event to Madagascar [30–32]. Yoder et al. [30] assessed the four potential models that could explain the appearance(s) of the euplerid ancestor on Madagascar: Gondwanan vicariance; landbridge connections; and ‘sweepstakes’ over-water dispersal in a single event (i.e., ‘rafting’) or in multiple steps via island-hopping. They concluded that available data are most consistent with the common ancestor of a monophyletic Eupleridae arriving in Madagascar ∼24-18 million years ago via a single over-water dispersal from African ancestry. Eupleridae consists of three major clades: Galidiinae, Euplerinae, and the fossa (*Cryptoprocta ferox*) (Fig. 1). Galidiines are comprised of five species of vontsira. Their superficial resemblance to mongooses (Herpestidae) have led to their “Malagasy mongoose” alias and even to their classification as herpestids [26,27]. Galidiines exhibit diverse ecologies such as in diets that range from insectivory, similarly found in herpestids, to more carnivorous diets in *Galidictis* species [22]. Euplerines superficially resemble civets (Viverridae), leading to them having been described as “Malagasy civets” and classified as viverrids [28]. Euplerines also exhibit diverse ecologies ranging from the omnivorous fanaloka (*Fossa fossana*) that feeds on a variety of invertebrate and vertebrate prey, similar to viverrids, to the Eastern falanouc (*Eupleres goudotii*) that specialize on soft-bodied invertebrates such as worms and slugs [22]. The last euplerid clade consists of the fossa, the largest and most carnivorous of the extant euplerids [33]. Its large-bodied civet-like and carnivorous cat-like resemblances have historically led its placement either within Viverridae or Felidae [28,29].

**Fig. 1.**
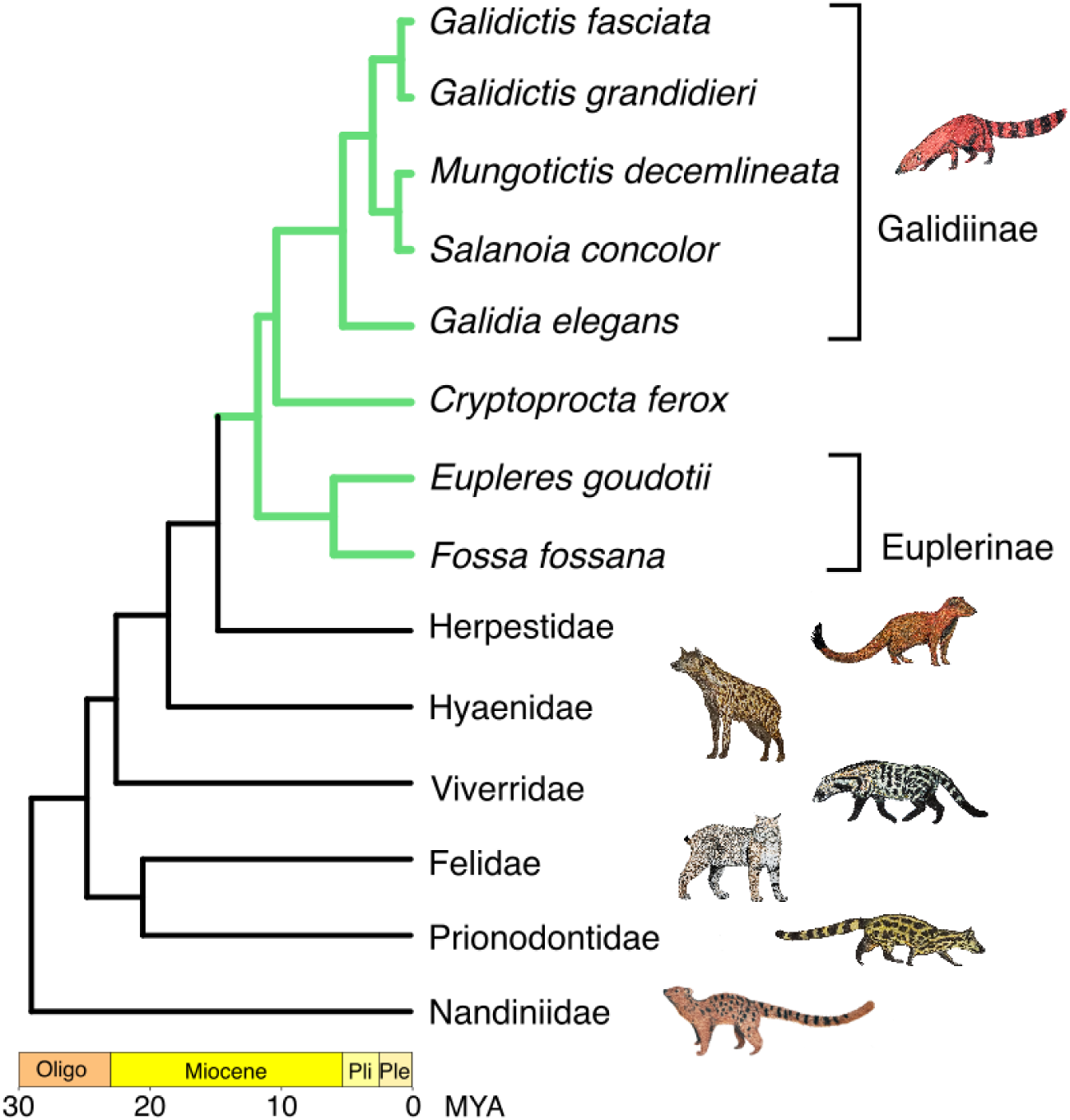
Phylogeny of Eupleridae and other extant feliform Carnivora families. Oligo = Oligocene; Pli = Pliocene; Ple = Pleistocene. Phylogeny based on [46].

Here, we investigated the morphological disparity among Malagasy euplerids and in comparison to taxa in other feliform carnivoran groups with which they have been historically classified. We used skull shape in our examination of phenotypic evolution because of its strong associations with dietary ecology across carnivorans [34–37]. Because of their superficial resemblances, we also investigated convergence among euplerid and feliform clades with which they have been classified historically. We predict that euplerids will exhibit greater disparity compared to other feliform clades and that the three euplerid clades—galidiines, euplerines, and the fossa—will converge towards herpestid-, viverrid-, and felid-like morphotypes in skull morphology, respectively. The observed phenotypic and ecological diversity found in euplerids suggest that they exhibited an adaptive radiation. Therefore, we also test if rates of their lineage diversification and phenotypic evolution are significantly accelerated relative to other clades as predicted under adaptive radiation. We predict that euplerids will exhibit shifts towards faster rates of lineage diversification and phenotypic evolution compared to other feliform clades.

## Methods

### Morphological data

We quantified the shapes of 110 crania across 57 feliforms (∼50% of species diversity) and the shapes of 315 mandibles across 90 feliforms (∼79% of species diversity) using 3D geometric morphometrics [38,39]. Our dataset encompasses all eight euplerid species (number of specimens per species ranged from 1–5). Whether *Eupleres major* and *Salanoia durrelli* are true species remains debated [40,41]. Shape data were collected from 3D scans of feliform skulls obtained from surface scanning with an EinScan Pro HD scanner at various museums, an Artec Spider structured light 3D scanner at the American Museum of Natural History, or from previously published work [35] (see Table S1 for list of specimens and museums). All specimens were fully mature, determined by the closure of exoccipital–basioccipital and basisphenoid– basioccipital sutures on the cranium and full tooth eruption. Although carnivorans exhibit sexual dimorphism [42,43], a combination of females, males, and sex unknown individuals was used because just one sex cannot be used without compromising the inclusion of as many euplerid species as possible. We used 35 landmarks and seven curves with 134 semi-landmarks for the cranium, and 21 landmarks and four curves with 24 semi-landmarks for the mandible (Fig. S1). Landmarks were digitized using Slicer, and curves were digitized by oversampling semi-landmarks in Slicer. Landmarks were superimposed by Generalized Procrustes analysis, and semi-landmarks on the curves were allowed to slide along their tangent vectors until their positions minimized bending energy [39,44]. As part of the superimposition procedure, bilaterally homologous landmarks and semi-landmarks were reflected across the median plane and averaged using the geomorph function bilat.symmetry. All Procrustes superimpositions were performed in the R package geomorph v4.0.6 [45]. We used centroid size as our metric of cranial and mandibular size.

### Craniomandibular shape analyses

We performed all analyses under a phylogenetic framework using a phylogeny of mammals generated by [46] and pruned to include only feliform carnivorans. All analyses were performed in R v4.1.1 (R Core Team 2021). Because allometry has been shown to facilitate or constrain skull shape evolution [47], we first tested for evolutionary allometry on cranial and mandibular shape by performing a phylogenetic Procrustes regression [48] with a random residual permutation procedure (1,000 iterations) using the geomorph function procD.pgls. Both cranial shape (SS = 0.022, MS = 0.022, R^2^ = 0.20, F = 14.127, Z = 3.45, P < 0.001) and mandibular shape (SS = 0.044, MS = 0.044, R^2^ = 0.25, F = 29.360, Z = 5.07, P = 0.001) exhibited significant evolutionary allometry. We, therefore, extracted allometry-free shape as the shape residuals from the phylogenetic Procrustes regressions and used them in all subsequent analyses.

#### Craniomandibular shape disparity

We visualized the phylomorphospace of cranial and mandibular shape by performing principal component analyses (PCA) using the geomorph function gm.prcomp. We then examined if cranial and mandibular disparity (i.e., Procrustes variance) differed among the feliform clades using the geomorph functions procD.plgs and morphol.disparity.

#### Testing for convergence towards different feliform morphotypes

Euplerids are often cited to superficially resemble and were historically classified together with other feliforms, specifically galidiines with mongooses (Herpestidae), euplerines with civets (Viverridae), and the fossa with cats (Felidae) or with civets. Therefore, we tested whether there was statistical evidence for convergence in these various euplerid-feliform comparisons. Here, we define convergence as lineages evolving to be more similar to one another than were their ancestors [49–51]. We assessed convergence using the Ct-measure in the R package convevol [50,51], which tests whether the geometric distance between focal lineages in phylomorphospace are shorter than the distances between ancestral nodes at a specific time point. Second, we used a frequency-based measure of convergence (C_5_), which determines whether the number of lineages that crosses into a particular region of morphospace is greater than expected. We conducted 100 simulations under Brownian motion to assess statistical significance of Ct and C_5_ measures. Together, these measures provide a more complete test of our definition of convergence.

We also tested whether each of the three groups exhibited shared adaptive zones in skull shapes by fitting five multivariate evolutionary models [52,53] on the first three PCs of the cranial shape dataset (accounting for 74.44% of total cranial shape variation) and mandibular shape dataset (87.4% of total mandibular shape variation) using the R package mvMORPH version 1.1.7 [54] to incorporate covariances between axes. These models included single-rate Brownian motion model (mvBM1), single-peak Ornstein–Uhlenbeck model (mvOU1), and three two-peak Ornstein–Uhlenbeck models (mvOUM). Each of these mvOUM models designated two optima, one optimum for the focal comparison and a second optimum for the remaining feliforms. All models were fit across 250 mapped trees to account for uncertainty in phylogenetic topology and stochastic character maps of group designations. Models were assessed with small sample corrected Akaike weights (AICcW), and all models with a ΔAICc < 2 were considered equally best-fitting models. Support for a mvOUM model as a best-fitting model would suggest a shared adaptive zone between the focal clades of interest. These results, however, also would provide more nuanced interpretations of the convergent evolution because they are unable to distinguish between potential convergence or conservatism [50,51].

#### Craniomandibular shape evolutionary rates

We investigated rates of cranial shape and mandibular shape evolution using two approaches. First, we tested if evolutionary rates in cranial shape and mandibular shape differed among the feliform clades using the geomorph function compare.evol.rates. Second, we examined branch-specific evolutionary rates and rate shifts of cranial shape and mandibular shape using a variable rates model under a reversible-jump Markov Chain Monte Carlo (rjMCMC) framework in BayesTraits v4.1.1 (http://www.evolution.rdg.ac.uk/). To reduce the dimensions of the shape data for BayesTraits analysis, we used only the phylogenetic principal component scores (pPC) scores that represent >95% of the shape variation in the cranium (first 18 pPCs) and mandible (first 11 pPCs). We ran two independent chains, each with 200,000,000 iterations sampled every 20,000 iterations. After the first 25,000,000 iterations were discarded as burn-in, we assessed convergence of the two chains by checking the trace plots and then the effective sample size (ESS) with Gelman and Rubin’s diagnostics in the R package coda v0.19-4. We plotted rates shifts and branch-specific rates across the feliform phylogeny and constructed density plots to compare evolutionary rates among feliform families using custom functions from [55]. A caveat to these analyses is that, outside of Eupleridae, we have more limited representation of ∼50% and ∼79% of species diversity of the remaining feliform clades in our cranial and mandibular shape datasets, respectively. Nevertheless, although identification of significant rate shifts potentially could be influenced by sampling in those other feliform clades, they are not the focus of this study.

### Rates of lineage diversification

We estimated speciation and extinction rates through time and across the feliform phylogeny using two methods, Bayesian Analysis of Macroevolutionary Mixtures (BAMM) [56] and the Lineage Specific Birth–Death–Shift (LSBDS) model [57]. [46]BAMM applies a reversible jump Markov Chain Monte Carlo (RJMCMC) to explore candidate models of lineage diversification as well as quantify heterogeneity in evolutionary rates. We performed two independent BAMM runs of five million generations on feliform phylogeny, sampling every 1,000 generations and with priors chosen using the R package BAMMtools v2.0 [58]. We assessed the convergence of each BAMM run using the R package BAMMtools v2.1.11 (Rabosky et al. 2014). The LSBDS model samples rate regimes from a prior distribution, discretized into a fixed number of rate categories. We implemented the LSBDS model using seven categories (one for each of the seven extant feliform families) for speciation and extinction, and ran two MCMC chains for 5,000 generations in RevBayes [59]. We merged the posteriors, retaining the last 4,000 generations from the MCMC in the R package RevGadgets [60]. Convergence for both methods was assessed by checking that the ESS values for all model parameters in the log files were greater than 200, using the R package coda [61].

## Results

### Craniomandibular shape morphospace and disparity

PCs 1–3 explain 74.44% of the cranial shape variation across feliforms (Fig. 2). PC 1 distinguishes hypercarnivorous felids and hyaenids (negative PC 1) from the remaining, less carnivorous feliforms including euplerids (positive PC 1). Positive PC 1 is characterized by lateral narrowing of the cranium at the zygomatic arches, expansion of the braincase through widening of the nuchal crests, and reduction of the sagittal crest resulting in dorsoventral flattening. The diversity of euplerid cranial shapes is best captured by PC 2, as euplerid species occupy the full range of PC 2. Positive PC 2 describes shortening of the rostrum and broadening of the cranium at the zygomatic arches and nuchal crests. Galidiines are associated with positive PC 2 and are encompassed within the range of variation of herpestids (and felids), whereas euplerines are associated with negative PC 2 with *Fossa fossana* falling within the range of viverrids (and hyaenids) but *Eupleres goudotii* distinct from all other feliforms. In PC2, *Cryptoprocta* is intermediate between galidiines and euplerines (closer to the former), and falls within the broad range of values of felids. In the total PC1/2 cranial morphospace, galidiines cluster with herpestids, euplerines are closest to and overlap with viverrids, and *Cryptoprocta* is distinct from other feliforms. Euplerids show little variation along cranial PC 3, falling entirely within the range of felids and overlapping with the lower range of viverrids and upper range of herpestids. All other non-monotypic feliform families show a greater range of variance than and/or do not overlap with euplerids in PC3. Positive PC 3 describes slight elongation and ventrodorsal flexion of the rostrum and slight broadening of the zygomatic arches. Ancestral node reconstruction suggests that euplerids exhibit a cranial shape between viverrids and herpestids (Fig. 3). Overall, euplerids exhibit significantly greater cranial shape disparity (Procrustes variance = 0.0078) compared to felids (0.0047, P = 0.026) and viverrids (0.0033, P = 0.004), but not hyaenids (0.0066, P = 0.595) or herpestids (0.0061, P = 0.299).

**Fig. 2.**
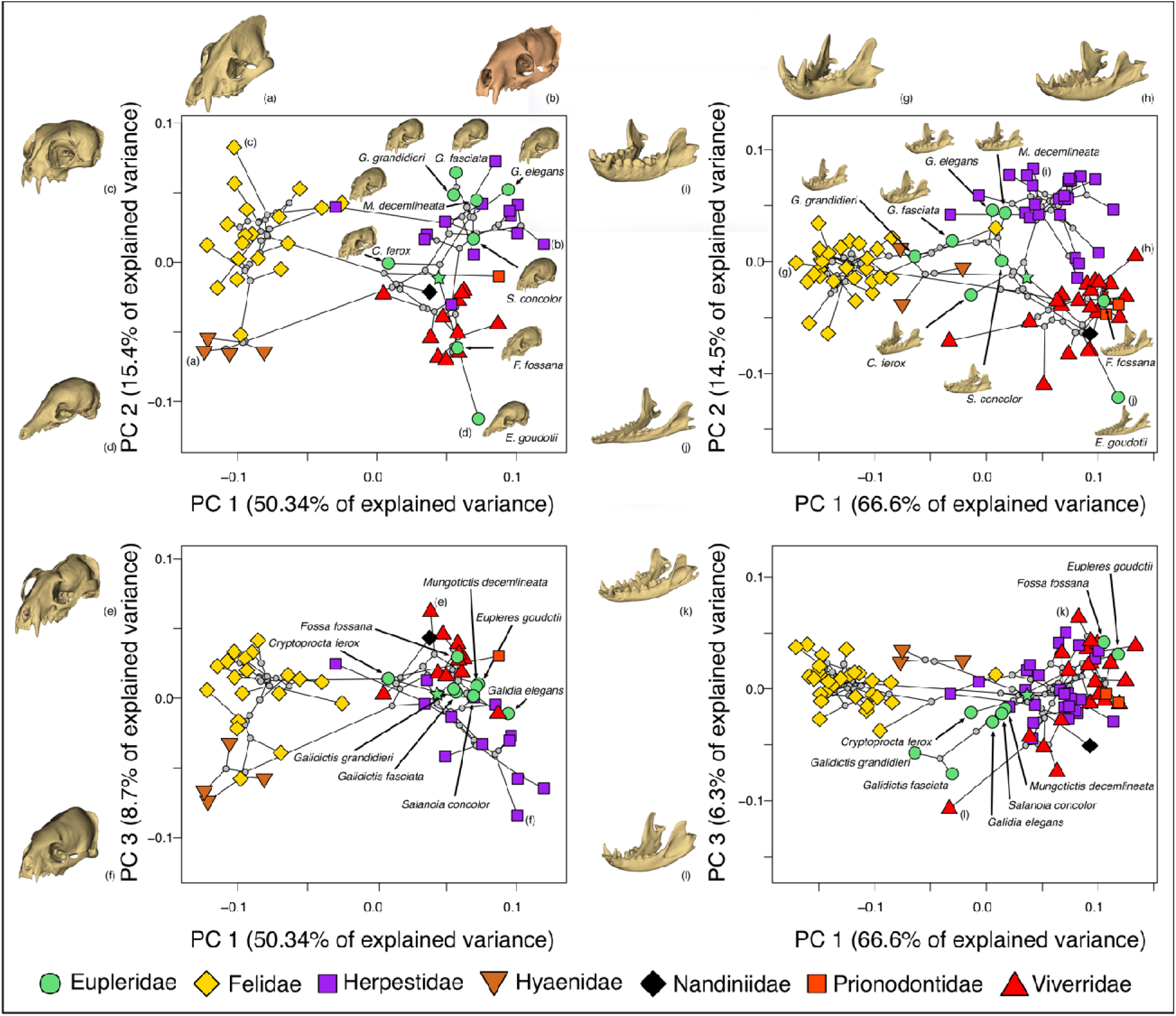
Morphospace of allometry-free cranial (left) and mandibular (right) shape defined by principal component (PC) axes 1–3. Taxa illustrated for the cranial morphospace: (a) Brown hyena (*Hyaena brunnea* [Hyaenidae]), –PC1; (b) Javan mongoose (*Urva javanicus* [Herpestidae]), +PC1; (c) Pallas’s cat (*Otocolobus manul* [Felidae]), +PC2; (d) Eastern falanouc (*Eupleres goudotii* [Eupleridae]), –PC2; (e) Asian palm civet (*Paradoxurus hermaphroditus* [Eupleridae]), +PC3; and (f) Egyptian mongoose (*Herpestes ichneumon* [Herpestidae]), –PC3. Taxa illustrated for the mandibular morphospace: (g) tiger (*Panthera tigris* [Felidae]), –PC1; (h) small Indian civet (*Viverricula indica* [Viverridae]), +PC1; (i) Ethiopian dwarf mongoose (*Helogale hirtula* [Herpestidae]), +PC2; (j) Eastern falanouc (*Eupleres goudotii* [Eupleridae]), – PC2; (k) Large Indian civet (*Viverra zibetha* [Viverridae]), +PC3; and (l) Binturong (*Arctictis binturong* [Viverridae]), –PC3. Grey circles indicate estimated ancestral shapes at nodes, and the green star indicates estimated ancestral shape of the ancestral euplerid node.

**Fig. 3.**
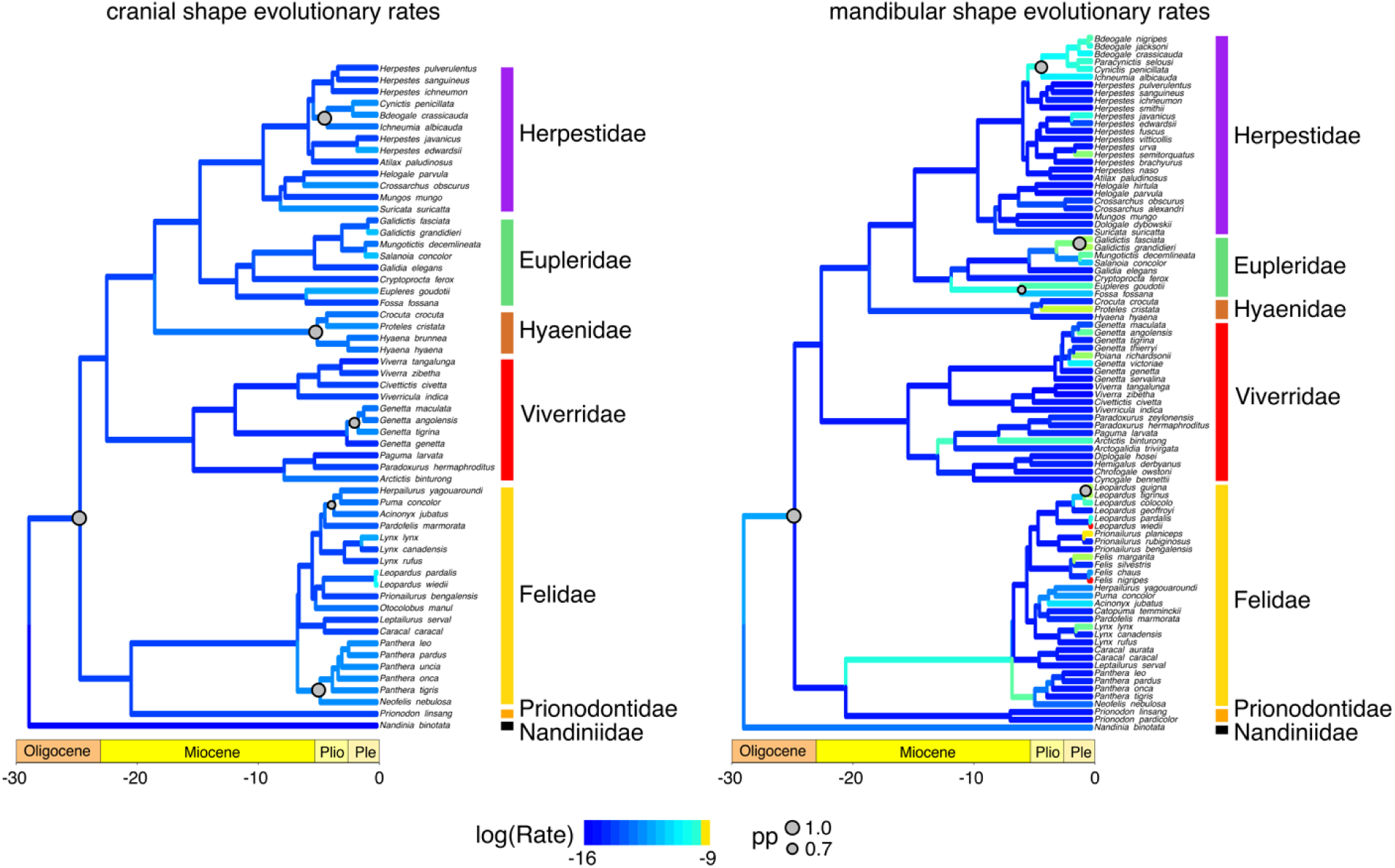
Phylorate plots of cranial and mandibular shape evolutionary rates across feliform Carnivora. Warmer colors indicate faster rates and cooler colors indicate slower rates. Significant shifts in evolutionary rate are shown as grey circles with sizes proportional to the posterior probability. All rates were log-transformed.

Euplerids occupy greater regions of mandibular morphospace when compared to cranial morphospace. PCs 1–3 explain 87.4% of the mandibular shape variation across feliforms (Fig. 2). Like in the crania, PC 1 of mandibular shape largely distinguishes hypercarnivorous felids, and to a lesser extent hyaenids (both with negative PC 1 values), from the remaining, less carnivorous feliforms (positive PC 1), except for overlap with hyaenids of two species of *Galidictis* (galidiine euplerids) and one herpestid. In mandibular PC1, euplerines fall within the range of viverrids and a few herpestids at the extreme positive end of their range of; galidiines have a wide range of mandibular PC1 values, overlapping with herpestids at the most negative end of their range of variation, hyaenids, and one species each of felid and viverrid; *Cryptoprocta* is very close to hyaenids in PC1 value. Positive PC 1 describes dorsoventral mandibular flexure and elongation, decrease in coronoid height, and lateral compression of the mandibular body. However, unlike in the crania, euplerids exhibit greater variation along PC 1 than in any other feliform family (although viverrids are similar in range of variation, due to one extreme outlier, the binturong [*Arctictis binturong*]), and spans negative and positive PC 1.

Euplerids also approach both the negative and positive ends of variation in both PCs 2 and 3, with *Eupleres goudotii* again having the most extreme negative value for PC2 of any feliform (as for the cranial PCA). Positive PC 2 describes anteroposterior shortening of the mandibular body, increases in coronoid height, and lateral broadening of the mandibular body; positive PC 3 describes anteroposterior elongation of the mandibular body, dorsoventral compression of the mandibular body, and reduction in coronoid height. Ancestral node reconstruction suggests that euplerids exhibit a cranial shape between viverrids and herpestids, with closer resemblance to the latter (Fig. 3). Euplerids exhibit significantly greater mandibular shape disparity (Procrustes variance = 0.0100) compared to felids (0.0035, P = 0.001), viverrids (0.0054, P = 0.020), and herpestids (0.0040, P = 0.003), but not hyaenids (0.0050, P = 0.692).

### Convergence towards different feliform morphotypes

We did not find statistical support for the distance-based measure of convergence between galidiines and herpestids (cranial Ct_1_ = -0.19, P = 0.96; mandibular Ct_1_ = -0.03, P = 0.39), between euplerines and viverrids (cranial Ct_1_ = -0.24, P = 0.84; mandibular Ct_1_ = -0.25, P = 0.84), between the fossa (*Cryptoprocta ferox*) and felids (cranial Ct_1_ = 0.05, P = 0.08; mandibular Ct_1_ = -0.05, P = 0.48) or between the fossa and viverrids (cranial Ct_1_ = -0.31, P = 0.98; mandibular Ct_1_ = -0.17, P = 0.46). Similarly, the frequency-based measure of convergence indicated that the number of transitions into regions of phylomorphospace occupied by herpestids, viverrids, and felids was not significantly greater than expected under Brownian motion (all P > 0.380).

Using evolutionary modeling, we found evidence of shared adaptive zones in cranial shape between galidiines and herpestids (best model = mvOUM, AICcW = 0.98) and some evidence of shared adaptive zones in cranial shape between euplerines and viverrids (best models = mvOUM [AICcW = 0.52] and mvBM [AICcW = 0.47; ΔAICc = 0.20]) and between the fossa and felids (best models = mvBM [AICcW = 0.66] and mvOUM [AICcW = 0.32; ΔAICc = 1.47]) (Table S2). In contrast, we found no evidence of shared adaptive zones in mandibular shape in all three comparisons (best model for all = mvOU1 [AICcW > 081]; Table S2).

### Rates of craniomandibular shape evolution

Rates of cranial shape evolution in euplerids (ln σ^2^_multi_ = -12.86) do not significantly differ from most feliform clades (felid ln σ^2^_multi_ = -12.94, P = 0.592; hyaenid ln σ^2^_multi_ = -12.47, P = 0.100; herpestid ln σ^2^_multi_ = -13.14, P = 0.105), except for viverrids (ln σ^2^_multi_ = -13.72, P = 0.001), in which euplerids exhibited significantly faster rates of cranial shape evolution. Although we found six significant rate shifts (posterior probability (pp) > 0.70) across feliforms, none of these occurred within euplerids (Fig. 3). Instead, rate shifts occurred towards or within the branches of the other major feliform clades including Hyaenidae, Felidae (pantherines and the *Puma* lineage), Viverridae (crownward *Genetta* species), Herpestidae (subclade of herpestines), and the overall feliform clade excluding Nandiniidae (Fig. 3).

Euplerids exhibited greater rate heterogeneity in evolution of mandibular shape compared to cranial shape. Euplerids exhibited significantly faster rates of mandibular shape evolution (ln σ^2^_multi_ = -12.29) compared to viverrids (ln σ^2^_multi_ = -13.08, P = 0.001) and herpestids (ln σ^2^_multi_ = -12.79, P = 0.001) but significantly slower rates compared to felids (ln σ^2^_multi_ = -11.80, P = 0.023). There was no difference in mandibular shape evolutionary rates between euplerids and hyaenids (ln σ^2^_multi_ = -12.23, P = 0.889). Furthermore, we found that two of the five significant rate shifts in mandibular shape (pp > 0.70) occurred within euplerid clades: *Galidictis* species and euplerines (Fig. 3). The remaining rate shifts occurred within the same subclade of herpestines that exhibited a rate shift in cranial shape evolution, within *Leopardus* cats, and for the overall feliform clade excluding Nandiniidae (Fig. 3).

### Diversification rate analyses

Euplerids do not exhibit distinct rates of diversification, as both BAMM and the LSBDS model are united in revealing that there are no increased species diversification rates on the branches leading towards Eupleridae compared to the remaining feliforms (Fig. 4; Fig. S2).

**Fig. 4.**
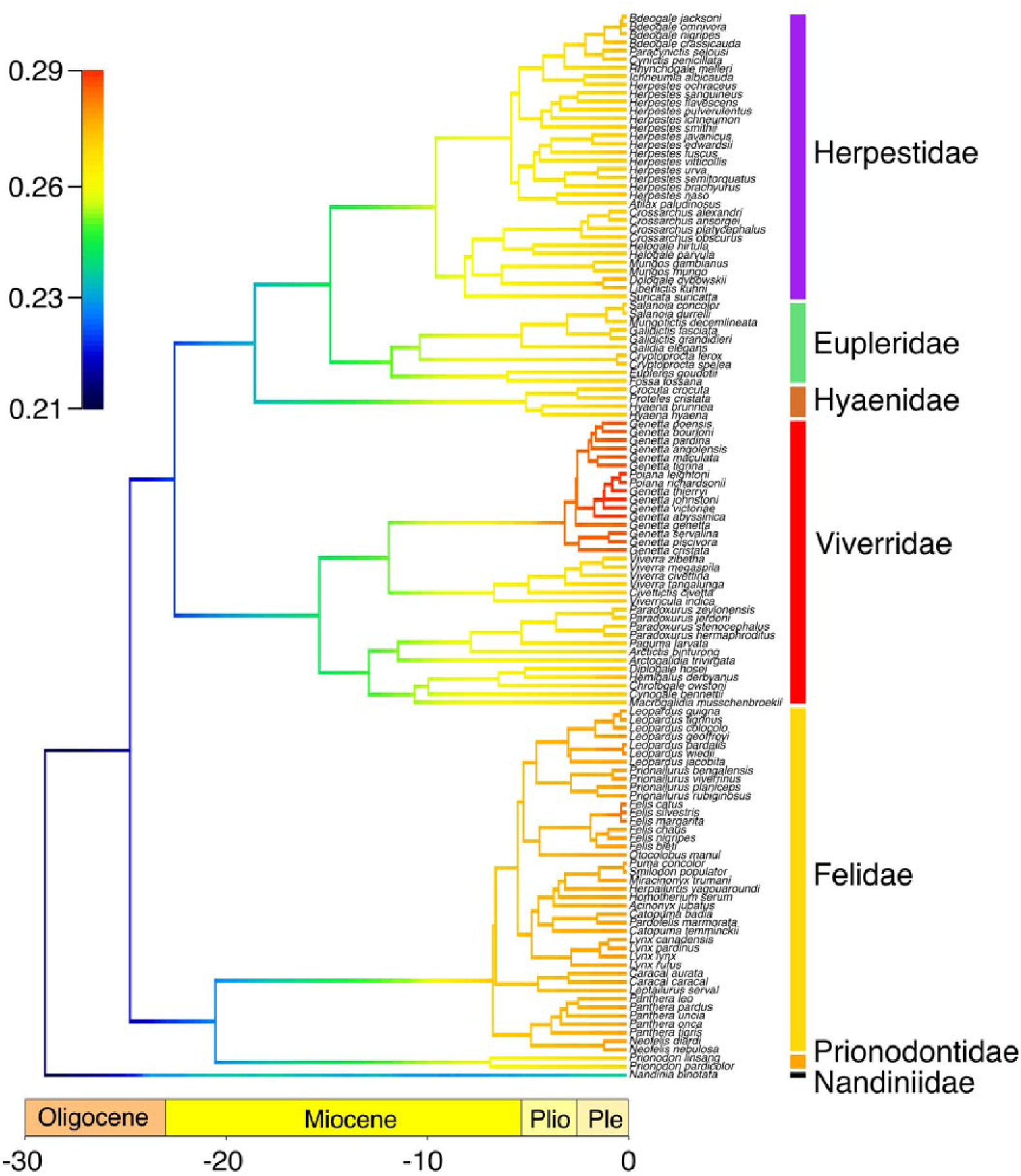
Phylorate plot of lineage diversification rates across feliform Carnivora using BAMM. Colors at each point in time along the branches of the phylorate plot denote instantaneous rate of diversification. Warmer colors indicate faster rates and cooler colors indicate slower rates. There are no significant shifts in diversification rates across the phylogeny. Analyses of lineage diversification rates using LSBDS resulted in similar results (Fig. S2).

Furthermore, we found no significant shifts in diversification rate within feliforms, albeit there are slightly elevated rates on the branches leading towards to the viverrid subfamily Genettinae and felids (Fig. 4; Fig. S2).

## Discussion

### Skull diversity in Malagasy carnivorans

Euplerids exhibit greater disparity in their skulls compared to most feliform families, for both cranial and mandibular shapes, and particularly so in their mandibles. Euplerids overlap with all extant feliform families in mandibular shape morphospace, whereas they only overlap with non-hyercarnivorous families (i.e., Herpestidae, Nandiniidae, Prionodontidae, and Viverridae) in cranial shape morphospace (Fig. 2). The only euplerid to approach hypercarnivorous feliforms (Felidae, Hyaenidae) in both cranial and mandibular morphospace is the fossa (*Cryptoprocta ferox*), the most carnivorous of the euplerids [22]. The fossa exhibits cranial and mandibular morphologies that appear intermediate between the average euplerid and average felid (Fig. 5A, 6A). Compared to all other euplerids, the fossa exhibits relatively broader zygomatic arches and more pronounced sagittal and nuchal crests in the cranium (Fig. 5A) and relatively broader coronoid processes in the mandible (Fig. 6A). These adaptations suggest that the fossa exhibits larger temporalis and masseter jaw muscles for increased biting ability [62–65] and are thus better adapted for their hypercarnivorous diets [22]. There is some evidence that the fossa and felids share a similar adaptive zone in cranial shape (ΔAIC = 1.47; Table S2), but there is no statistical support for convergence in skull shape between the fossa and felids. These results suggest that cranial adaptations towards hypercarnivory in the fossa do not quite reach the felid morphotype (Fig. 5A, 6A) because the fossa and felids share an adaptive zone characterized by broad adaptive slopes rather than distinct adaptive peaks with steep slopes [24,51,66,67].

**Fig. 5.**
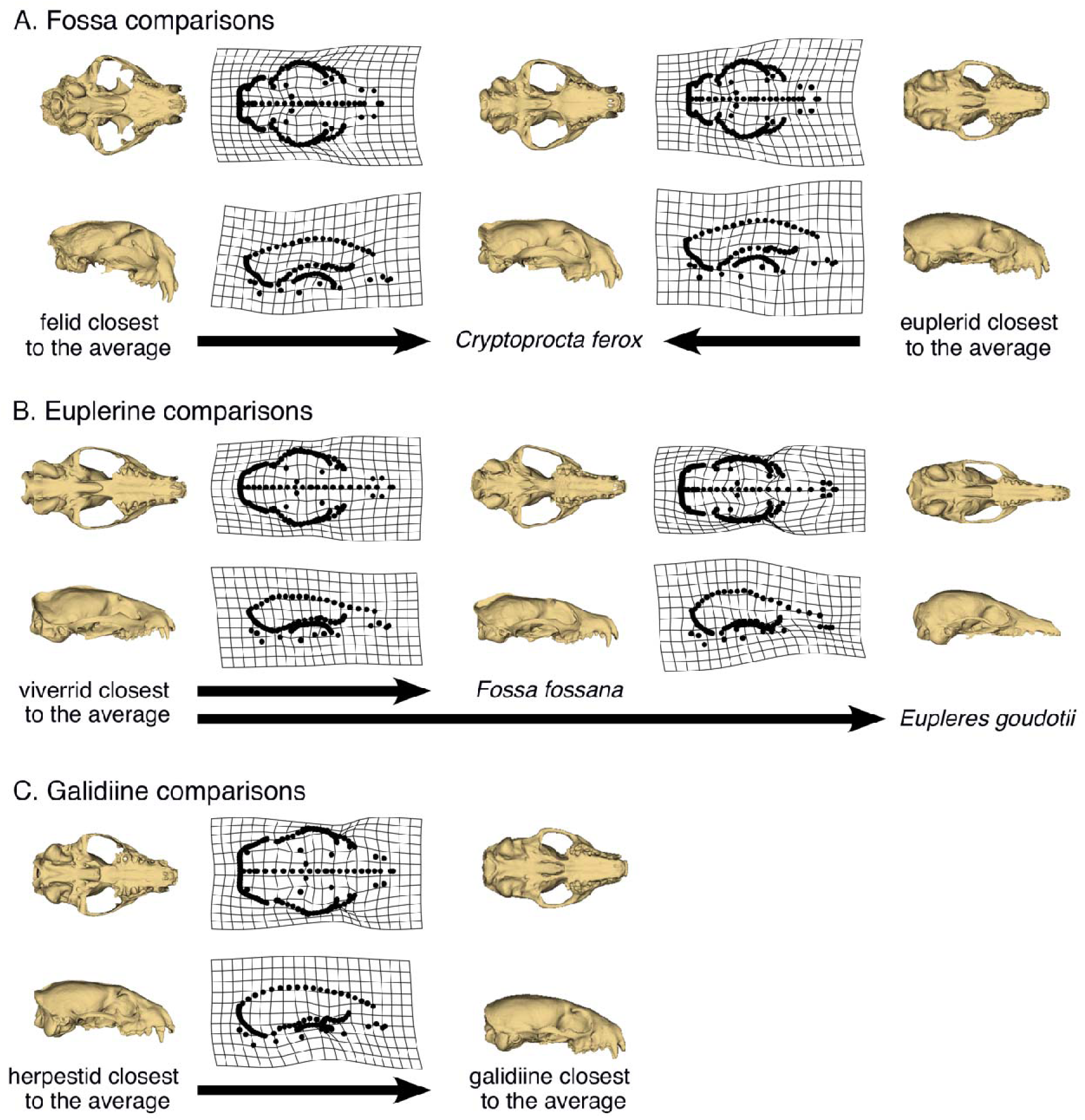
Thin-plate spline (TPS) deformation grids depicting cranial shape differences between euplerid clades and other feliform families. A) Cranial shape of the fossa (*Crytoprocta ferox*) appears intermediate between felids and other euplerids. B) Cranial shape of euplerines: the fanaloka (*Fossa fossana*) resembles viverrids whereas the Eastern falanouc (*Eupleres goudotii*) exhibits a more specialized morphology. C) Cranial shapes of most galidiines resemble herpestids.

**Fig. 6.**
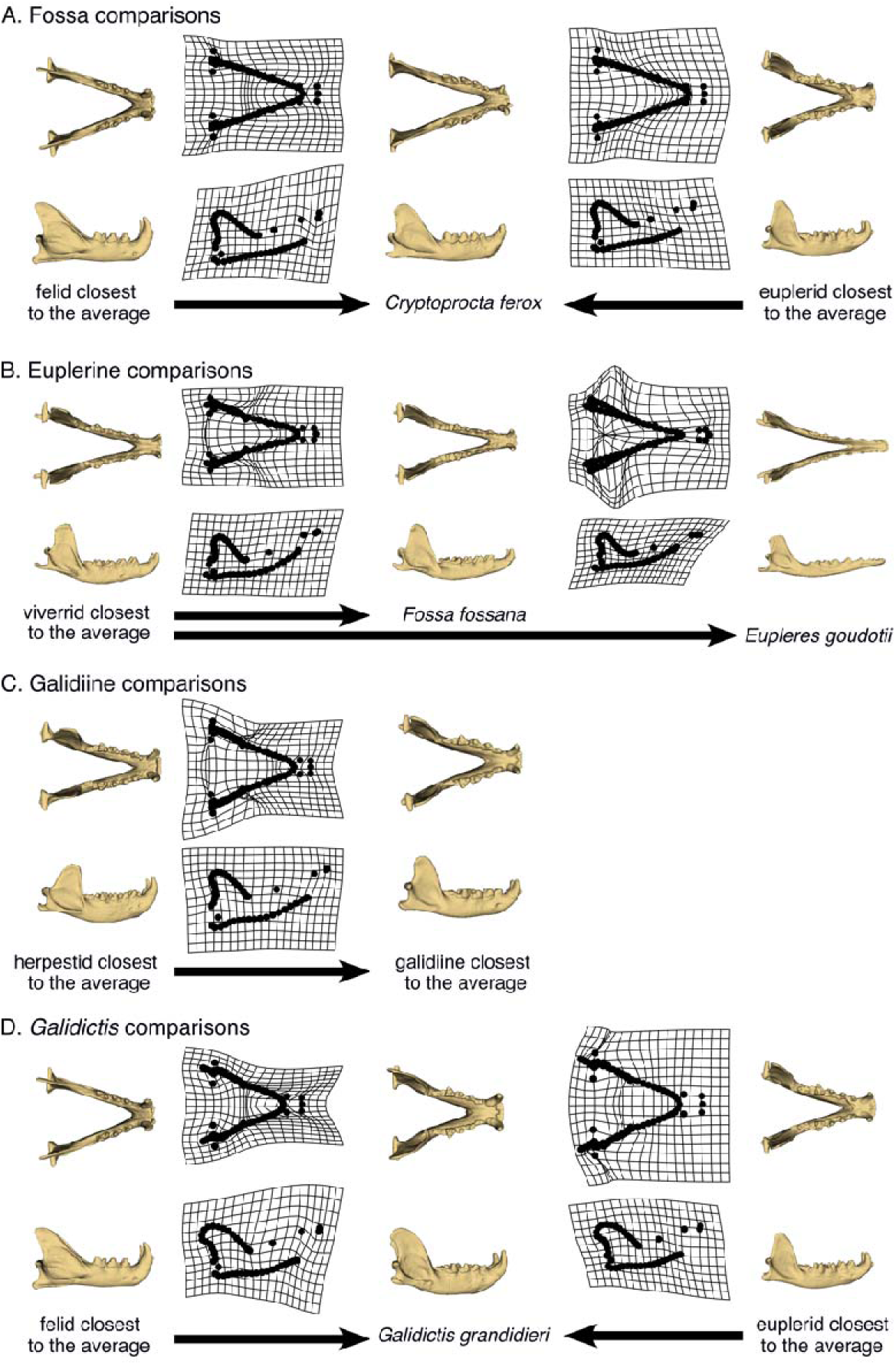
Thin-plate spline (TPS) deformation grids depicting mandibular shape differences euplerid clades and other feliform families. A) Mandibular shape of the fossa (*Crytoprocta ferox*) appears intermediate between felids and euplerids. B) Mandibular shape of euplerines: the fanaloka (*Fossa fossana*) resemble viverrids whereas the Eastern falanouc (*Eupleres goudotii*) exhibits a more specialized morphology. C) Mandibular shapes of most galidiines resemble herpestids, although the mandibular shape (D) of Grandidier’s vontsira (*Galidictis grandidieri*) appears intermediate between felids and other galidiines.

Euplerines occupy overlapping regions of cranial and mandibular morphospace with viverrids, the family to which they have been compared or classified with historically (Fig. 2). The fanaloka (*Fossa fossana*) exhibits only slight differences in skull shape (i.e., narrower and more elongate crania and mandibles) compared to the average viverrid, which is consistent with their similar diets of invertebrate and vertebrate prey, whereas the falanouc (*Eupleres goudotii*) exhibits a more specialized skull that includes relatively elongate rostrum and mandible and less pronounced zygomatic arches, nuchal crest, sagittal crests, coronoid processes compared to the average viverrid (Fig. 5B) and all other feliforms. These adaptations in the skull, along with reduced dentition, enable the falanouc to specialize in feeding on soft-bodied invertebrates such as worms and slugs [22]. We found evidence that euplerines and viverrids share an adaptive zone in cranial shape but not mandibular shape (Table S2), but neither skull component provided statistical support for convergence between the two groups. Therefore, like for the fossa and felids, our results suggest that euplerines and viverrids may share an adaptive zone with broad adaptive slopes instead of exhibiting convergence characterized by distinct similar adaptive peaks with steep slopes.

Galidiines exhibit distinct patterns of cranial and mandibular shape distribution in morphospace. Galidiines occupy overlapping regions of cranial morphospace with herpestids (Fig. 2), and there appear to be few cranial shape differences between the galidiine species closest to the average cranial shape of all galidiines and the herpestid species closest to the average cranial shape of all herpestids (Fig. 5C). Evolutionary modeling also strongly supports a shared adaptive zone in cranial shape between galidiines and herpestids (Table S2) but no statistical support for convergence. However, galidiines approach regions of mandibular morphospace occupied by hypercarnivorous feliforms (Fig. 2) and exhibit mandibular morphologies that appear intermediate between herpestids and felids (Fig. 6C, D): galidiine mandibles tend to be relatively broader and more robust compared to the herpestid species closest to the average mandibular shape of all herpestids, but are not as broad or robust as the felid species closest to the average mandibular shape of all felids (Fig. 6C, D). Of the galidiines, *Galidictis* species appear most similar to felids in mandibular shape, providing morphological evidence that carnivory likely is an important component of their diet. Previous work surmised that in addition to insects, *Galidictis* feed on rodents, reptiles, amphibians, and even small lemurs; however, their dietary ecologies remain poorly known, as they are some of the rarest carnivorans in the world [22]. That *Galidictis* exhibits a rate shift towards faster mandibular evolution (Fig. 3) leads us to postulate that shifts towards more carnivorous diets may be due to more relatively recent (< 5 million years ago) selection. The distinct patterns between the cranium and mandible in galidiines follow an overall trend of decoupled modes of evolution between the cranium and mandible across carnivorans, where cranial evolution tends to follow clade-based shifts whereas mandibular evolution is linked instead to broad dietary regimes [35,68]. Our results suggest that galidiines may have retained cranial morphology from euplerid-herpestid ancestors through conservatism rather than convergence whereas the more evolutionary labile mandible adapted towards a more carnivorous diet than the typical insectivorous diets found in herpestids, most euplerids, and most likely their most recent common ancestors (of the euplerid + herpestid clade).

### Are Malagasy carnivorans an adaptive radiation?

Like many endemic Malagasy clades, euplerids are often considered to represent an adaptive radiation after dispersal to a long-isolated island. We found that euplerids show extreme ranges of skull shape variation and many distinct cranial and mandibular shapes in phylomorphospace and geometric morphometric analyses. Contrary to predictions of adaptive radiation, we found that euplerids do not exhibit exceptional lineage diversification and have only marginally faster rates of mandibular shape evolution and, to a lesser extent, cranial shape evolution compared to other feliform families (Fig. 3, 4). Thus, it is tempting to postulate that euplerids may have evolved via an “adaptive non-radiation” [8], where ecomorphological divergence within a clade fails to produce rapid species diversification. Rapid phenotypic evolution without rapid taxonomic diversification rates have been found in a variety of vertebrate clades [69–71] including other carnivorans such as mustelids [72] as well as other endemic Malagasy vertebrates such as mantellid frogs [8]. Those studies cited possible factors that may mask a signal of rapid diversification [8,69–72] including the absence of fossil data in diversification analyses, which may have erased signatures of early rapid diversification [73–75]; continental radiations, in which many clades are simply unable to rapidly radiate spatially across entire continents as soon as they arise [70,72,76–78]; and clade age, where younger clades diversify faster than older clades [79]. These factors also may influence our analyses of euplerid diversification. First, euplerids are not an exceptionally young clade, having diverged from herpestids ∼24–18 million years ago [30] with a clade age for extant species of ∼15–14 million years old [46]. Despite their long evolutionary history, euplerids are extremely poorly represented in the fossil record, known only from Holocene subfossil remains of the extinct giant fossa (*Cryptoprocta spelea*) [80]. Therefore, unaccounted cladogenesis of extinct euplerids may lead to underestimation of taxonomic diversification rates and ecological diversity in euplerids. Second, unaccounted speciation of modern euplerids also may lead to underestimation of diversification rates. At least two euplerids—*Eupleres goudotii* and *Salanoia concolor*—are thought, though with uncertainty, to each consist of two species (i.e., *E. goudotii* and *E. major* and *S. concolor* and *S. durrelli*) [40,41,81]; if valid, this would lead to increased diversification rates, albeit at more crownward tips of the tree. Lastly, euplerid diversification is perhaps better characterized as a continental radiation rather than as an insular radiation. Researchers often suggest that Madagascar, as the 4^th^ largest island globally, resembles a continent more than an island, with a geological record that spans more than three billion years and several vicariance and dispersal events over the past ∼88-90 million years, after its final separation from other major Gondwanan landmasses [82–84]. Therefore, the diversification of Malagasy clades with long residence times in Madagascar may be better characterized as multiple bursts of speciation events throughout their evolutionary history [19,85] rather than as a single early burst of rapid diversification that has been found in some other insular clades [9–11]. That diversification rates do not differ statistically between euplerids and continental feliforms support the hypothesis that the diversification dynamics of other endemic Malagasy clades follow a pattern more similar to diversification on a continent than on an island [8]. Therefore, while found only on Madagascar, there is no indication that insular processes spurred a rapid burst of taxonomic diversification of euplerids, as often has been hypothesized.

## Conclusion

Euplerids exhibit great disparity in cranial and mandibular morphology that corresponds to their wide dietary diversity ranging from worm-eating falanoucs to lemur-eating fossas. We document for the first time that their skull morphologies, particularly their mandibles, show significantly greater shape disparity compared to other feliform clades, and this single small clade of Malagasy endemic carnivorans captures a large amount of the morphological variation that is found across the entire clade of feliforms. We also found evidence that the three euplerid clades—galidiines, euplerines, and the fossa—exhibited shared adaptive zones in cranial shape morphology with herpestids, viverrids, and felids, respectively. We did not find statistical evidence, however, that these three euplerid clades—galidiines, euplerines, and the fossa— converged towards herpestid-, viverrid-, and felid-morphotypes in skull morphology, respectively, as had been implicit in classifications or directly hypothesized historically. Despite demonstrating great skull disparity, we found that euplerids do not clearly exhibit an adaptive radiation, as we found no evidence of rapid lineage diversification rates and only marginally faster rates of mandibular shape evolution compared to other feliform families. This work supports an increasing number of studies demonstrating that adaptive radiations in endemic Malagasy clades may not as common as previously thought [8,19] and diversification dynamics on Madagascar may resemble those found on continents rather than on islands.

## Acknowledgements

We thank C. Merrill for scanning of euplerid specimens at AMNH, and R. Andrade Luna, E. Blackwell, F. Levy, and J. Padro for helping scan and landmark specimens. We are grateful to the staff of and collections at the American Museum of Natural History, California Academy of Sciences, Field Museum of Natural History, Natural History Museum of Los Angeles County, Museum of Vertebrate Zoology, Natural History Museum London, National Museum of Natural History, and Burke Museum of Natural History and Culture.

## Ethics

This work did not require ethical approval from a human subject or animal welfare committee.

## Data accessibility

All data and original code are available from the Dryad Digital Repository: https://datadryad.org/stash/dataset/doi:10.5061/dryad.x95x69psj

## Declaration of AI use

We have not used AI-assisted technologies in creating this article.

