## Supplementary Materials for "Skull evolution and lineage diversification in endemic Malagasy carnivorans"

Electronic supplementary material for

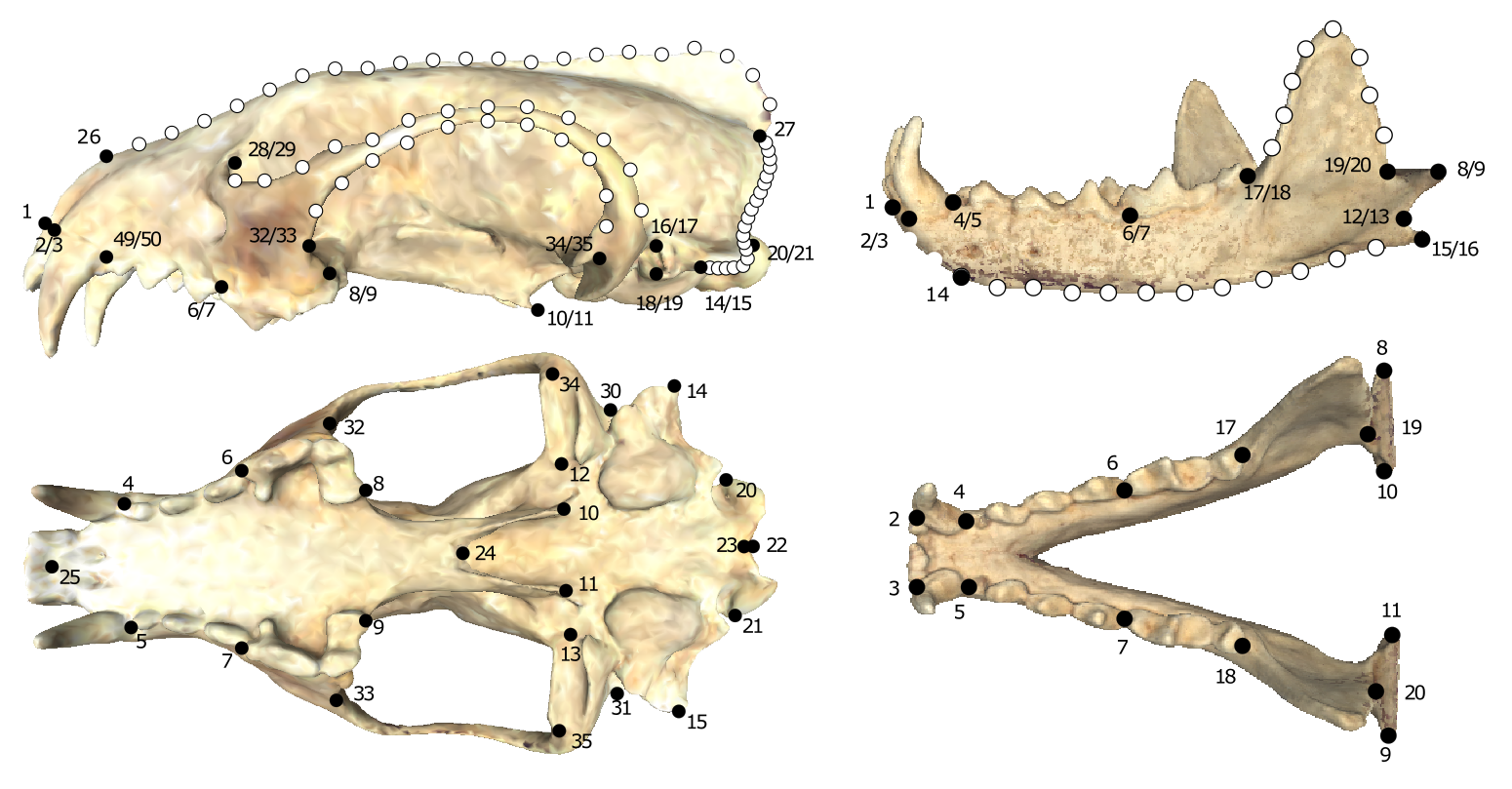

Fig. S1. Geometric morphometric landmarks and curves. Cranium: L1) Anteriormost point of premaxilla. L2 & L3) Anteriormost point on canine alveolus. L4 & L5) Anteriormost point on premolar alveolus. L6 & L7) Anteriomost point on the first molar alveolus. L8 & L 9) Posteriormost point on the last molar alveolus. L10 & L11) Ventralmost point on pterygoid hamulus. L12 & L13) Medialmost margin of the mandibular fossa. L14 & L15) Ventralmost point on mastoid process. L16 & L17) Dorsalmost point on the external edge of the auditory meatus. L18 & L19) Ventralmost point on the external edge of the auditory meatus. L20 & L21) Lateralmost point on occipital condyle. L22) Dorsal border of foramen magnum. L23) Ventral border of foramen magnum. L24) Posteriormost point on midline of palate. L25) Anteriormost point on midline of the complete palate. L26) Anteriormost point on the midline of the nasals. L27) Posteriormost point on the intersection of the lambdoidal and sagittal crests. L28 & L29) Anteriormost point on the inflection of the orbit. L30 & L31) Posteriormost point on the intersection of the zygomatic arch and braincase. L32 & L33) Ventralmost point of the insertion of the zygomatic arch on the maxilla. L34 & L35) Lateralmost point on the margin of the mandibular fossa. Curve 1 from L26 to L27, along the dorsal midline of the cranium. Curves 2 & 3 from L28/29 to L30/31, along the dorsal profile of the zygomatic arch. Curves 4 & 5 from L32/33 to L34/35, along the ventral profile of the zygomatic arch. Curves 6 & 7 from L14/L15 to L27 along the occipital ridge. *Mandible:* L1) Anteriormost point on the mandibular symphysis. L2 & L3) Anteriormost point on canine alveolus. L4 & L5) Anteriormost point on premolar alveolus. L6 & L7) Anteriormost point of first molar alveolus. L8 & L9) Lateralmost point on mandibular condyle. L10 & L11) Medialmost point on mandibular condyle. L12 & L13) Inflection point on the posterior profile between the mandibular condyle and the angular process. L14) Ventralmost point on the mandibular symphysis. L15 & L16) Posteriormost point on the angular process. L17 & L18) Posteriormost point on the last molar alveolus. L19 & L20) Anterior inflection point on the articular surface, midpoint between L8/9 and L10/11. Curves 1 & 2 from L17/18 to L19/20, along the dorsal profile of the coronoid process. Curves 3 & 4 from L14 to L15-16, along the ventral profile of the ramus and angular process.

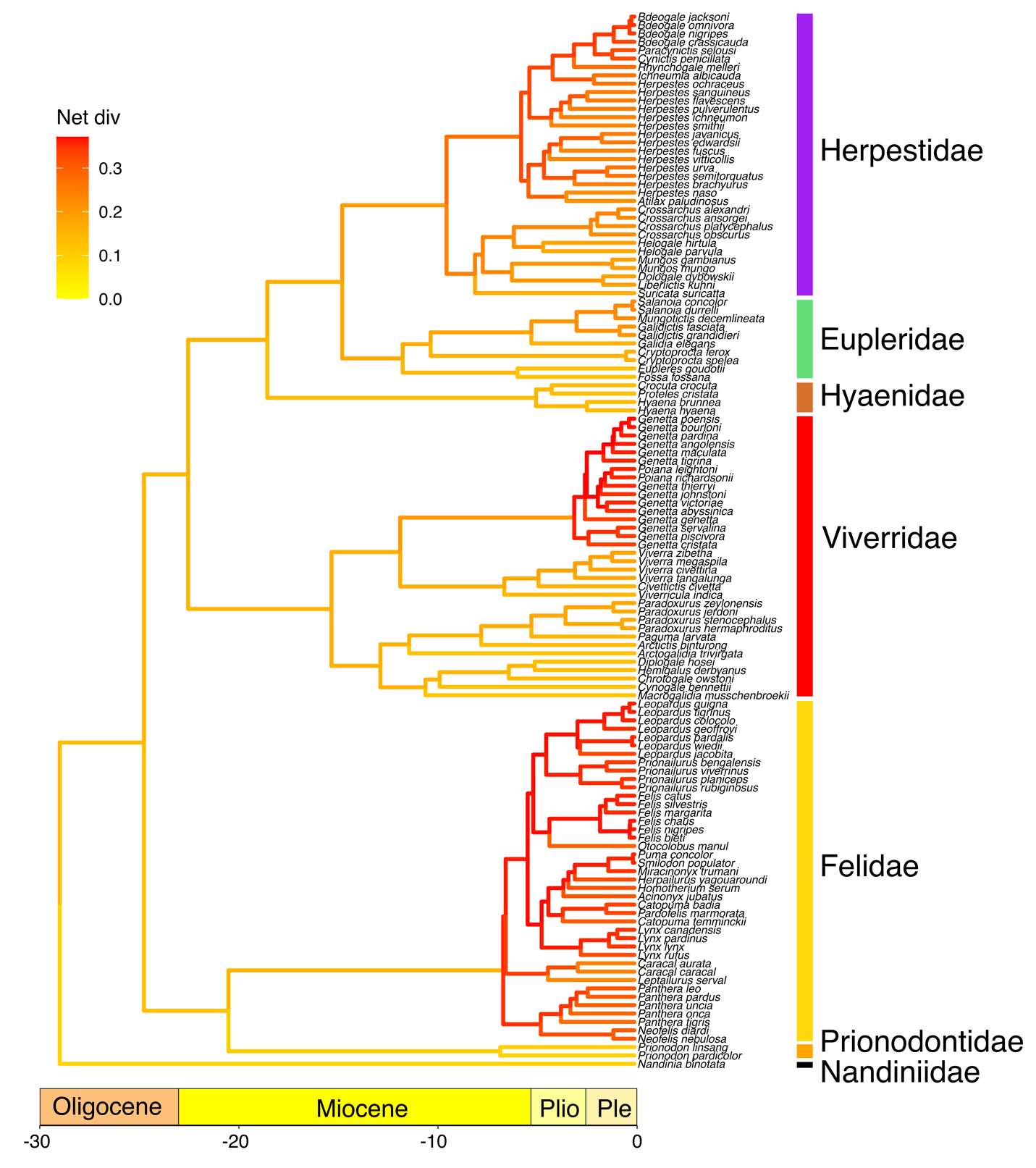

Fig. S2. Phylorate plot of lineage diversiﬁcation rates across feliform Carnivora using the Lineage Specific Birth–Death–Shift (LSBDS) model. Colors at each point in time along the branches of the phylorate plot denote instantaneous rate of diversiﬁcation. Redder colors indicate faster rates and yellower colors indicate slower rates. There are no signiﬁcant shifts in diversiﬁcation rates across the phylogeny. Analyses of lineage diversification rates using BAMM resulted in similar results (Fig. 4).

**Table S1**. List of specimens used in the study. AHR = Adam Heartstone-Rose personal collection; AMNH = American Museum of Natural History; CAS = California Academy of Sciences; DMNS = Denver Museum of Nature & Science; FMNH = Field Museum; LACM = Natural History Museum of Los Angeles County; MNHN = Muséum national d'Histoire naturelle; MVZ = Museum of Vertebrate Zoology; NHMUK = Natural History Museum in London; SMNS = Stuttgart State Museum of Natural History; USNM = U.S. National Museum of Natural History; UWBM = Burke Museum; ZMB = Museum für Naturkunde. -ms indicates specimen was obtained from Morphosource.

| A. Crania | |  |  |  |
| --- | --- | --- | --- | --- |
|  | family | species | sex | catalog |
|  | 01Nandiniidae | *Nandinia_binotata* | M | lacm53740 |
|  | 02Prionodontidae | *Prionodon_linsang* | U | USNM539732 |
|  | 03Felidae | *Acinonyx_jubatus* | F | FMNH127834 |
|  | 03Felidae | *Acinonyx_jubatus* | M | FMNH29635 |
|  | 03Felidae | *Caracal_caracal* | U | AHR |
|  | 03Felidae | *Felis_lybica* | F | LACM14474 |
|  | 03Felidae | *Felis_lybica* | M | LACM14480 |
|  | 03Felidae | *Herpailurus_yagouaroundi* | F | dmb5816-ms |
|  | 03Felidae | *Herpailurus_yagouaroundi* | M | zmb5815-ms |
|  | 03Felidae | *Leopardus_pardalis* | M | FMNH34339 |
|  | 03Felidae | *Leopardus_wiedii* | F | LACM29326 |
|  | 03Felidae | *Leptailurus_serval* | U | AHR |
|  | 03Felidae | *Lynx_canadensis* | F | UWBM33180 |
|  | 03Felidae | *Lynx_canadensis* | M | UWBM82238 |
|  | 03Felidae | *Lynx_lynx* | F | smns001870-ms |
|  | 03Felidae | *Lynx_rufus* | F | LACM10118 |
|  | 03Felidae | *Lynx_rufus* | F | LACM15254 |
|  | 03Felidae | *Lynx_rufus* | M | LACM10115 |
|  | 03Felidae | *Lynx_rufus* | M | UWBM31939 |
|  | 03Felidae | *Neofelis_nebulosa* | M | USNM28214 |
|  | 03Felidae | *Otocolobus_manul* | M | UWBM35449 |
|  | 03Felidae | *Panthera_leo* | M | UWBM33191 |
|  | 03Felidae | *Panthera_leo* | U | MVZ117849 |
|  | 03Felidae | *Panthera_onca* | U | AHR |
|  | 03Felidae | *Panthera_onca* | U | AHR |
|  | 03Felidae | *Panthera_pardus* | M | LACM11704 |
|  | 03Felidae | *Panthera_tigris* | F | MVZ189634-MS |
|  | 03Felidae | *Panthera_uncia* | F | UWBM34194 |
|  | 03Felidae | *Panthera_uncia* | M | LACM54708 |
|  | 03Felidae | *Pardofelis_marmorata* | U | LACM469 |
|  | 03Felidae | *Prionailurus_bengalensis* | M | cas34385 |
|  | 03Felidae | *Puma_concolor* | F | LACM85440 |
|  | 03Felidae | *Puma_concolor* | F | UWBM51188 |
|  | 03Felidae | *Puma_concolor* | M | LACM87430 |
|  | 04Viverridae | *Arctictis_binturong* | U | AHR |
|  | 04Viverridae | *Arctictis_binturong* | F | USNM259101 |
|  | 04Viverridae | *Civettictis_civetta* | F | dmnh17711-ms |
|  | 04Viverridae | *Civettictis_civetta* | M | dnmnh1190-ms |
|  | 04Viverridae | *Genetta_angolensis* | F | MVZ118449 |
|  | 04Viverridae | *Genetta_genetta* | U | mnhn-zm-mo1997-450-MS |
|  | 04Viverridae | *Genetta_maculata* | F | smns004416-ms |
|  | 04Viverridae | *Genetta_maculata* | M | smns004425-ms |
|  | 04Viverridae | *Genetta_tigrina* | F | UWBM35891 |
|  | 04Viverridae | *Paguma_larvata* | F | UWBM73282 |
|  | 04Viverridae | *Paguma_larvata* | U | UWBM73281 |
|  | 04Viverridae | *Paradoxurus_hermaphroditus* | F | cas21131 |
|  | 04Viverridae | *Paradoxurus_hermaphroditus* | M | cas34386 |
|  | 04Viverridae | *Paradoxurus_hermaphroditus* | U | mnhn-zm-ac-a3448-MS |
|  | 04Viverridae | *Paradoxurus_hermaphroditus* | U | mnhn-zm-ac-a3448-MS |
|  | 04Viverridae | *Viverra_tangalunga* | F | cas9465 |
|  | 04Viverridae | *Viverra_tangalunga* | F | smns002257-ms |
|  | 04Viverridae | *Viverra_zibetha* | F | lacm8220 |
|  | 04Viverridae | *Viverra_zibetha* | u | smns002507-ms |
|  | 04Viverridae | *Viverricula_indica* | M | uwbm |
|  | 04Viverridae | *Viverricula_indica* | u | smns021594-ms |
|  | 05Hyaenidae | *Crocuta_crocuta* | F | USNM164506 |
|  | 05Hyaenidae | *Crocuta_crocuta* | M | USNM181527 |
|  | 05Hyaenidae | *Hyaena_hyaena* | M | USNM182034 |
|  | 05Hyaenidae | *Hyaena_brunnea* | M | FMNH345874 |
|  | 05Hyaenidae | *Hyaena_brunnea* | U | MVZ117842 |
|  | 05Hyaenidae | *Proteles_cristata* | F | USNM368497 |
|  | 05Hyaenidae | *Proteles_cristata* | U | j050607t02-MS |
|  | 06Eupleridae | *Cryptoprocta_ferox* | M | AMNH188213 |
|  | 06Eupleridae | *Cryptoprocta_ferox* | M | amnh188213-ms |
|  | 06Eupleridae | *Cryptoprocta_ferox* | M | NHMUK32-7-19-2 |
|  | 06Eupleridae | *Eupleres_goudotii* | F | AMNH100462 |
|  | 06Eupleridae | *Eupleres_goudotii* | M | NHMUK35-1-8-308 |
|  | 06Eupleridae | *Eupleres_goudotii* | M | FMNH30492 |
|  | 06Eupleridae | *Fossa_fossana* | F | AMNH188209 |
|  | 06Eupleridae | *Fossa_fossana* | F | FMNH85196 |
|  | 06Eupleridae | *Fossa_fossana* | M | AMNH188210 |
|  | 06Eupleridae | *Fossa_fossana* | M | NHMUK35-1-8-293 |
|  | 06Eupleridae | *Galidia_elegans* | F | AMNH100465 |
|  | 06Eupleridae | *Galidia_elegans* | F | FMNH162110 |
|  | 06Eupleridae | *Galidia_elegans* | M | AMNH100464 |
|  | 06Eupleridae | *Galidia_elegans* | M | AMNH100466 |
|  | 06Eupleridae | *Galidia_elegans* | M | FMNH167984 |
|  | 06Eupleridae | *Galidictis_fasciata* | F | NHMUK1938-11-16-5 |
|  | 06Eupleridae | *Galidictis_fasciata* | M | FMNH162111 |
|  | 06Eupleridae | *Galidictis_fasciata* | U | AMNH100479 |
|  | 06Eupleridae | *Galidictis_grandidieri* | F | FMNH173158 |
|  | 06Eupleridae | *Mungotictis_decemlineata* | F | FMNH184087 |
|  | 06Eupleridae | *Mungotictis_decemlineata* | M | FMNH176128 |
|  | 06Eupleridae | *Salanoia_concolor* | M | NHMUK1938-11-16-8 |
|  | 06Eupleridae | *Salanoia_concolor* | M | NHMUK80-469 |
|  | 07Herpestidae | *Atilax_paludinosus* | F | dmnh9019-ms |
|  | 07Herpestidae | *Atilax_paludinosus* | M | dmnh20426-ms |
|  | 07Herpestidae | *Bdeogale_crassicauda* | F | lacm56752 |
|  | 07Herpestidae | *Bdeogale_crassicauda* | M | lacm42940 |
|  | 07Herpestidae | *Crossarchus_obscurus* | U | AHR |
|  | 07Herpestidae | *Cynictis_penicillata* | F | MVZ117834 |
|  | 07Herpestidae | *Cynictis_penicillata* | F | MVZ117836 |
|  | 07Herpestidae | *Herpestes_pulverulentus* | F | MVZ117829 |
|  | 07Herpestidae | *Herpestes_pulverulentus* | M | LACM30761 |
|  | 07Herpestidae | *Helogale_parvula* | F | lacm56754 |
|  | 07Herpestidae | *Helogale_parvula* | M | lacm59638 |
|  | 07Herpestidae | *Herpestes_edwardsii* | U | UWBM41669 |
|  | 07Herpestidae | *Herpestes_ichneumon* | F | lacm45745 |
|  | 07Herpestidae | *Herpestes_ichneumon* | M | lacm30317 |
|  | 07Herpestidae | *Herpestes_javanicus* | U | AHR |
|  | 07Herpestidae | *Herpestes_javanicus* | M | amnh101655-MS |
|  | 07Herpestidae | *Herpestes_sanguineus* | F | MVZ117824 |
|  | 07Herpestidae | *Herpestes_sanguineus* | F | MVZ118456 |
|  | 07Herpestidae | *Herpestes_sanguineus* | M | MVZ117827 |
|  | 07Herpestidae | *Herpestes_sanguineus* | M | MVZ118459 |
|  | 07Herpestidae | *Ichneumia_albicauda* | M | LACM40138 |
|  | 07Herpestidae | *Mungos_mungo* | F | MVZ118466 |
|  | 07Herpestidae | *Mungos_mungo* | M | cas12223 |
|  | 07Herpestidae | *Suricata_suricatta* | F | MVZ117823 |
|  | 07Herpestidae | *Suricata_suricatta* | F | MVZ118450 |
| B. Mandibles | |  |  |  |
|  | family | *species* | sex | catalog |
|  | 01Nandiniidae | *Nandinia_binotata* | F | AMNH51474 |
|  | 01Nandiniidae | *Nandinia_binotata* | F | AMNH51475 |
|  | 01Nandiniidae | *Nandinia_binotata* | M | lacm53740 |
|  | 01Nandiniidae | *Nandinia_binotata* | M | usnm450440 |
|  | 02Prionodontidae | *Prionodon_linsang* | F | FMNH88300 |
|  | 02Prionodontidae | *Prionodon_linsang* | F | USNM395048 |
|  | 02Prionodontidae | *Prionodon_linsang* | M | USNM303036 |
|  | 02Prionodontidae | *Prionodon_linsang* | M | usnm539732 |
|  | 02Prionodontidae | *Prionodon_pardicolor* | F | FMNH35493 |
|  | 02Prionodontidae | *Prionodon_pardicolor* | M | FMNH35464 |
|  | 02Prionodontidae | *Prionodon_pardicolor* | M | FMNH39175 |
|  | 03Felidae | *Acinonyx_jubatus* | F | AMNH119655 |
|  | 03Felidae | *Acinonyx_jubatus* | F | AMNH119657 |
|  | 03Felidae | *Acinonyx_jubatus* | M | AMNH27897 |
|  | 03Felidae | *Acinonyx_jubatus* | M | fmnh29635 |
|  | 03Felidae | *Caracal_aurata* | F | AMNH89441 |
|  | 03Felidae | *Caracal_aurata* | M | AMNH51994 |
|  | 03Felidae | *Caracal_aurata* | M | NRMA585866 |
|  | 03Felidae | *Caracal_caracal* | F | AMNH187788 |
|  | 03Felidae | *Caracal_caracal* | M | MNHN-ZM-AC-1932-3229 |
|  | 03Felidae | *Catopuma_temminckii* | F | MNHN-ZO-AC-1939-2152 |
|  | 03Felidae | *Catopuma_temminckii* | M | FMNH89919 |
|  | 03Felidae | *Felis_chaus* | F | AMNH54528 |
|  | 03Felidae | *Felis_chaus* | F | FMNH103995 |
|  | 03Felidae | *Felis_chaus* | M | lacm8224 |
|  | 03Felidae | *Felis_chaus* | M | LACM8224 |
|  | 03Felidae | *Felis_lybica* | F | FMNH106749 |
|  | 03Felidae | *Felis_lybica* | F | FMNH140228 |
|  | 03Felidae | *Felis_lybica* | M | FMNH95877 |
|  | 03Felidae | *Felis_lybica* | M | FMNH98325 |
|  | 03Felidae | *Felis_margarita* | F | FMNH107299 |
|  | 03Felidae | *Felis_nigripes* | F | USNM381275 |
|  | 03Felidae | *Felis_silvestris* | F | AMNH133972 |
|  | 03Felidae | *Felis_silvestris* | F | AMNH96294 |
|  | 03Felidae | *Felis_silvestris* | M | LACM45759 |
|  | 03Felidae | *Felis_silvestris* | M | LACM56696 |
|  | 03Felidae | *Herpailurus_yagouaroundi* | F | AMNH46538 |
|  | 03Felidae | *Herpailurus_yagouaroundi* | F | LACM61144 |
|  | 03Felidae | *Herpailurus_yagouaroundi* | M | ZMB21295 |
|  | 03Felidae | *Herpailurus_yagouaroundi* | M | ZMB5815 |
|  | 03Felidae | *Leopardus_colocolo* | F | FMNH68318 |
|  | 03Felidae | *Leopardus_colocolo* | F | MVZ114777 |
|  | 03Felidae | *Leopardus_colocolo* | M | AMNH76150 |
|  | 03Felidae | *Leopardus_colocolo* | M | FMNH52488 |
|  | 03Felidae | *Leopardus_geoffroyi* | F | AMNH205905 |
|  | 03Felidae | *Leopardus_geoffroyi* | F | AMNH205908 |
|  | 03Felidae | *Leopardus_geoffroyi* | M | MVZ145342 |
|  | 03Felidae | *Leopardus_geoffroyi* | M | NRMA592001 |
|  | 03Felidae | *Leopardus_guigna* | F | AMNH33288 |
|  | 03Felidae | *Leopardus_pardalis* | F | AMNH181995 |
|  | 03Felidae | *Leopardus_pardalis* | F | AMNH67708 |
|  | 03Felidae | *Leopardus_pardalis* | M | LACM92559 |
|  | 03Felidae | *Leopardus_pardalis* | M | MVZ4907 |
|  | 03Felidae | *Leopardus_tigrinus* | F | AMNH80396 |
|  | 03Felidae | *Leopardus_tigrinus* | F | FMNH70569 |
|  | 03Felidae | *Leopardus_tigrinus* | M | FMNH79923 |
|  | 03Felidae | *Leopardus_tigrinus* | M | FMNH85823 |
|  | 03Felidae | *Leopardus_wiedii* | F | AMNH148994 |
|  | 03Felidae | *Leopardus_wiedii* | F | AMNH29596 |
|  | 03Felidae | *Leopardus_wiedii* | M | MVZ132175 |
|  | 03Felidae | *Leopardus_wiedii* | M | MVZ132176 |
|  | 03Felidae | *Leptailurus_serval* | F | FMNH90022 |
|  | 03Felidae | *Leptailurus_serval* | F | FMNH93310 |
|  | 03Felidae | *Leptailurus_serval* | M | SMNS018897 |
|  | 03Felidae | *Leptailurus_serval* | M | SMNS018901 |
|  | 03Felidae | *Lynx_canadensis* | F | AMNH127754 |
|  | 03Felidae | *Lynx_canadensis* | F | MVZ4166 |
|  | 03Felidae | *Lynx_canadensis* | M | MVZ88576 |
|  | 03Felidae | *Lynx_canadensis* | M | UWBM80612 |
|  | 03Felidae | *Lynx_lynx* | F | SMNS001870 |
|  | 03Felidae | *Lynx_lynx* | F | SMNS018860 |
|  | 03Felidae | *Lynx_lynx* | M | NRM20025024 |
|  | 03Felidae | *Lynx_rufus* | F | lacm10118 |
|  | 03Felidae | *Lynx_rufus* | F | lacm15254 |
|  | 03Felidae | *Lynx_rufus* | M | UWBM81437 |
|  | 03Felidae | *Lynx_rufus* | M | UWBM84186 |
|  | 03Felidae | *Neofelis_nebulosa* | M | usnm28214 |
|  | 03Felidae | *Panthera_leo* | F | AMNH19181 |
|  | 03Felidae | *Panthera_leo* | F | AMNH30243 |
|  | 03Felidae | *Panthera_leo* | M | MVZ117855 |
|  | 03Felidae | *Panthera_leo* | M | MVZ124260 |
|  | 03Felidae | *Panthera_onca* | F | AMNH120998 |
|  | 03Felidae | *Panthera_onca* | F | AMNH135929 |
|  | 03Felidae | *Panthera_onca* | M | MVZ4900 |
|  | 03Felidae | *Panthera_onca* | M | ZMB56267 |
|  | 03Felidae | *Panthera_pardus* | F | AMNH169459 |
|  | 03Felidae | *Panthera_pardus* | F | AMNH54854 |
|  | 03Felidae | *Panthera_pardus* | M | SMNS018948 |
|  | 03Felidae | *Panthera_pardus* | M | SMNS018957 |
|  | 03Felidae | *Panthera_tigris* | F | AMNH113743 |
|  | 03Felidae | *Panthera_tigris* | F | FMNH31797 |
|  | 03Felidae | *Panthera_tigris* | M | FMNH31153 |
|  | 03Felidae | *Panthera_tigris* | M | MNHN-ZO-AC-1931-60 |
|  | 03Felidae | *Pardofelis_marmorata* | F | LACM713 |
|  | 03Felidae | *Pardofelis_marmorata* | M | AMNH102844 |
|  | 03Felidae | *Prionailurus_bengalensis* | F | AMNH58371 |
|  | 03Felidae | *Prionailurus_bengalensis* | F | AMNH59957 |
|  | 03Felidae | *Prionailurus_bengalensis* | M | FMNH39166 |
|  | 03Felidae | *Prionailurus_bengalensis* | M | FMNH65458 |
|  | 03Felidae | *Prionailurus_planiceps* | F | USNM144119 |
|  | 03Felidae | *Prionailurus_planiceps* | F | USNM145594 |
|  | 03Felidae | *Prionailurus_rubiginosus* | F | FMNH96333 |
|  | 03Felidae | *Prionailurus_rubiginosus* | M | FMNH95037 |
|  | 03Felidae | *Puma_concolor* | F | lacm85440 |
|  | 03Felidae | *Puma_concolor* | F | UWBM39058 |
|  | 03Felidae | *Puma_concolor* | M | UWBM51197 |
|  | 03Felidae | *Puma_concolor* | M | ZMB37691 |
|  | 04Viverridae | *Arctictis_binturong* | F | usnm259101 |
|  | 04Viverridae | *Arctogalidia_trivirgata* | F | AMNH107054 |
|  | 04Viverridae | *Arctogalidia_trivirgata* | M | USNM122919 |
|  | 04Viverridae | *Arctogalidia_trivirgata* | M | USNM311450 |
|  | 04Viverridae | *Chrotogale_owstoni* | F | MVZ186571 |
|  | 04Viverridae | *Civettictis_civetta* | F | AMNH55461 |
|  | 04Viverridae | *Civettictis_civetta* | M | NHMUK27-12-21-24 |
|  | 04Viverridae | *Civettictis_civetta* | M | NHMUK66-776 |
|  | 04Viverridae | *Cynogale_bennettii* | F | USNM145587 |
|  | 04Viverridae | *Cynogale_bennettii* | F | USNM145588 |
|  | 04Viverridae | *Cynogale_bennettii* | M | AMNH103993 |
|  | 04Viverridae | *Cynogale_bennettii* | M | NHMUK79-11-21-633 |
|  | 04Viverridae | *Genetta_angolensis* | F | AMNH80748 |
|  | 04Viverridae | *Genetta_angolensis* | F | AMNH89205 |
|  | 04Viverridae | *Genetta_genetta* | F | AMNH169072 |
|  | 04Viverridae | *Genetta_genetta* | F | AMNH169073 |
|  | 04Viverridae | *Genetta_genetta* | M | MVZ118443 |
|  | 04Viverridae | *Genetta_genetta* | M | MVZ118444 |
|  | 04Viverridae | *Genetta_maculata* | F | SMNS004412 |
|  | 04Viverridae | *Genetta_maculata* | F | SMNS004414 |
|  | 04Viverridae | *Genetta_maculata* | M | SMNS004426 |
|  | 04Viverridae | *Genetta_maculata* | M | SMNS004427 |
|  | 04Viverridae | *Genetta_servalina* | F | AMNH51566 |
|  | 04Viverridae | *Genetta_servalina* | F | AMNH51568 |
|  | 04Viverridae | *Genetta_servalina* | M | FMNH145228 |
|  | 04Viverridae | *Genetta_servalina* | M | FMNH145230 |
|  | 04Viverridae | *Genetta_thierryi* | F | LACM36694 |
|  | 04Viverridae | *Genetta_thierryi* | F | LACM53727 |
|  | 04Viverridae | *Genetta_thierryi* | M | LACM45750 |
|  | 04Viverridae | *Genetta_thierryi* | M | USNM450964 |
|  | 04Viverridae | *Genetta_tigrina* | F | FMNH196628 |
|  | 04Viverridae | *Genetta_tigrina* | F | USNM351943 |
|  | 04Viverridae | *Genetta_tigrina* | M | LACM53731 |
|  | 04Viverridae | *Genetta_tigrina* | M | USNM469854 |
|  | 04Viverridae | *Genetta_victoriae* | F | AMNH51411 |
|  | 04Viverridae | *Genetta_victoriae* | M | AMNH51414 |
|  | 04Viverridae | *Genetta_victoriae* | M | AMNH51426 |
|  | 04Viverridae | *Hemigalus_derbyanus* | F | FMNH8374 |
|  | 04Viverridae | *Hemigalus_derbyanus* | F | FMNH85109 |
|  | 04Viverridae | *Hemigalus_derbyanus* | M | USNM084424 |
|  | 04Viverridae | *Hemigalus_derbyanus* | M | USNM197239 |
|  | 04Viverridae | *Diplogale_hosei* | M | FMNH74275 |
|  | 04Viverridae | *Paguma_larvata* | F | USNM124278 |
|  | 04Viverridae | *Paguma_larvata* | F | USNM198058 |
|  | 04Viverridae | *Paguma_larvata* | M | USNM258863 |
|  | 04Viverridae | *Paguma_larvata* | M | USNM292907 |
|  | 04Viverridae | *Paradoxurus_hermaphroditus* | F | AMNH185315 |
|  | 04Viverridae | *Paradoxurus_hermaphroditus* | F | AMNH207583 |
|  | 04Viverridae | *Paradoxurus_hermaphroditus* | M | FMNH91250 |
|  | 04Viverridae | *Paradoxurus_hermaphroditus* | M | WAM17386 |
|  | 04Viverridae | *Paradoxurus_zeylonensis* | M | FMNH99409 |
|  | 04Viverridae | *Poiana_richardsonii* | M | AMNH51438 |
|  | 04Viverridae | *Viverra_tangalunga* | F | AMNH152881 |
|  | 04Viverridae | *Viverra_tangalunga* | F | AMNH152882 |
|  | 04Viverridae | *Viverra_tangalunga* | M | AMNH152883 |
|  | 04Viverridae | *Viverra_tangalunga* | M | AMNH207582 |
|  | 04Viverridae | *Viverra_zibetha* | F | AMNH163593 |
|  | 04Viverridae | *Viverra_zibetha* | F | AMNH54781 |
|  | 04Viverridae | *Viverra_zibetha* | M | FMNH104395 |
|  | 04Viverridae | *Viverra_zibetha* | M | FMNH114374 |
|  | 04Viverridae | *Viverricula_indica* | F | AMNH102459 |
|  | 04Viverridae | *Viverricula_indica* | F | AMNH102914 |
|  | 04Viverridae | *Viverricula_indica* | M | LACM56772 |
|  | 04Viverridae | *Viverricula_indica* | M | uwbm56775 |
|  | 05Hyaenidae | *Crocuta_crocuta* | F | AMNH52064 |
|  | 05Hyaenidae | *Crocuta_crocuta* | F | MVZ165169 |
|  | 05Hyaenidae | *Crocuta_crocuta* | M | MVZ165160 |
|  | 05Hyaenidae | *Crocuta_crocuta* | M | MVZ165179 |
|  | 05Hyaenidae | *Hyaena_hyaena* | F | AMNH54512 |
|  | 05Hyaenidae | *Hyaena_hyaena* | F | FMNH101948 |
|  | 05Hyaenidae | *Hyaena_hyaena* | M | LACM70163 |
|  | 05Hyaenidae | *Hyaena_hyaena* | M | usnm182034 |
|  | 05Hyaenidae | *Proteles_cristata* | F | AMNH169089 |
|  | 05Hyaenidae | *Proteles_cristata* | F | AMNH169445 |
|  | 05Hyaenidae | *Proteles_cristata* | M | AMNH169091 |
|  | 05Hyaenidae | *Proteles_cristata* | M | AMNH27768 |
|  | 06Eupleridae | *Cryptoprocta_ferox* | F | FMNH161707 |
|  | 06Eupleridae | *Cryptoprocta_ferox* | F | USNM112841 |
|  | 06Eupleridae | *Cryptoprocta_ferox* | M | AMNH188213 |
|  | 06Eupleridae | *Cryptoprocta_ferox* | M | FMNH5655 |
|  | 06Eupleridae | *Cryptoprocta_ferox* | M | MNHN-ZO-AC-2017-3240 |
|  | 06Eupleridae | *Cryptoprocta_ferox* | M | NHMUK1938-11-16-1 |
|  | 06Eupleridae | *Cryptoprocta_ferox* | M | NHMUK32-7-19-2 |
|  | 06Eupleridae | *Eupleres_goudotii* | F | NHMUK35-1-8-309 |
|  | 06Eupleridae | *Eupleres_goudotii* | M | FMNH30492 |
|  | 06Eupleridae | *Eupleres_goudotii* | M | NHMUK35-1-8-308 |
|  | 06Eupleridae | *Eupleres_goudotii* | M | NHMUK35-1-8-311 |
|  | 06Eupleridae | *Fossa_fossana* | F | AMNH188209 |
|  | 06Eupleridae | *Fossa_fossana* | F | FMNH85196 |
|  | 06Eupleridae | *Fossa_fossana* | F | NHMUK35-1-8-290 |
|  | 06Eupleridae | *Fossa_fossana* | F | NHMUK72-8-79-4 |
|  | 06Eupleridae | *Fossa_fossana* | M | AMNH100454 |
|  | 06Eupleridae | *Fossa_fossana* | M | NHMUK35-1-8-289 |
|  | 06Eupleridae | *Fossa_fossana* | M | NHMUK35-1-8-293 |
|  | 06Eupleridae | *Galidia_elegans* | F | AMNH100471 |
|  | 06Eupleridae | *Galidia_elegans* | F | AMNH100473 |
|  | 06Eupleridae | *Galidia_elegans* | F | FMNH162110 |
|  | 06Eupleridae | *Galidia_elegans* | F | FMNH173110 |
|  | 06Eupleridae | *Galidia_elegans* | F | FMNH195829 |
|  | 06Eupleridae | *Galidia_elegans* | M | FMNH156650 |
|  | 06Eupleridae | *Galidia_elegans* | M | FMNH156651 |
|  | 06Eupleridae | *Galidia_elegans* | M | FMNH161921 |
|  | 06Eupleridae | *Galidia_elegans* | M | FMNH161925 |
|  | 06Eupleridae | *Galidia_elegans* | M | FMNH167984 |
|  | 06Eupleridae | *Galidictis_fasciata* | F | FMNH156549 |
|  | 06Eupleridae | *Galidictis_fasciata* | F | FMNH178720 |
|  | 06Eupleridae | *Galidictis_fasciata* | F | NHMUK1938-11-16-5 |
|  | 06Eupleridae | *Galidictis_fasciata* | F | USNM449210 |
|  | 06Eupleridae | *Galidictis_fasciata* | M | FMNH156652 |
|  | 06Eupleridae | *Galidictis_fasciata* | M | FMNH162111 |
|  | 06Eupleridae | *Galidictis_fasciata* | M | NHMUK1938-11-16-2 |
|  | 06Eupleridae | *Galidictis_fasciata* | M | NHMUK1938-11-16-6 |
|  | 06Eupleridae | *Galidictis_grandidieri* | F | FMNH173158 |
|  | 06Eupleridae | *Mungotictis_decemlineata* | F | FMNH184087 |
|  | 06Eupleridae | *Mungotictis_decemlineata* | M | FMNH176128 |
|  | 06Eupleridae | *Salanoia_concolor* | F | NHMUK1938-11-16-11 |
|  | 06Eupleridae | *Salanoia_concolor* | F | NHMUK25-4-10-10 |
|  | 06Eupleridae | *Salanoia_concolor* | M | NHMUK1938-11-16-8 |
|  | 06Eupleridae | *Salanoia_concolor* | M | NHMUK80-469 |
|  | 07Herpestidae | *Atilax_paludinosus* | F | AMNH42058 |
|  | 07Herpestidae | *Atilax_paludinosus* | F | AMNH51636 |
|  | 07Herpestidae | *Atilax_paludinosus* | M | LACM53752 |
|  | 07Herpestidae | *Atilax_paludinosus* | M | LACM53753 |
|  | 07Herpestidae | *Bdeogale_crassicauda* | F | FMNH211260 |
|  | 07Herpestidae | *Bdeogale_crassicauda* | F | LACM42942 |
|  | 07Herpestidae | *Bdeogale_crassicauda* | M | lacm42940 |
|  | 07Herpestidae | *Bdeogale_crassicauda* | M | LACM42941 |
|  | 07Herpestidae | *Bdeogale_jacksoni* | F | AMNH36024 |
|  | 07Herpestidae | *Bdeogale_jacksoni* | F | AMNH36025 |
|  | 07Herpestidae | *Bdeogale_jacksoni* | M | AMNH35942 |
|  | 07Herpestidae | *Bdeogale_jacksoni* | M | AMNH36028 |
|  | 07Herpestidae | *Bdeogale_nigripes* | F | AMNH51586 |
|  | 07Herpestidae | *Bdeogale_nigripes* | F | FMNH85969 |
|  | 07Herpestidae | *Bdeogale_nigripes* | M | FMNH167685 |
|  | 07Herpestidae | *Crossarchus_alexandri* | F | AMNH51643 |
|  | 07Herpestidae | *Crossarchus_alexandri* | F | AMNH51649 |
|  | 07Herpestidae | *Crossarchus_alexandri* | M | AMNH51641 |
|  | 07Herpestidae | *Crossarchus_alexandri* | M | AMNH51661 |
|  | 07Herpestidae | *Cynictis_penicillata* | F | LACM56715 |
|  | 07Herpestidae | *Cynictis_penicillata* | F | LACM59634 |
|  | 07Herpestidae | *Cynictis_penicillata* | M | LACM41573 |
|  | 07Herpestidae | *Cynictis_penicillata* | M | LACM56712 |
|  | 07Herpestidae | *Dologale_dybowskii* | M | AMNH51018 |
|  | 07Herpestidae | *Herpestes_pulverulentus* | F | AMNH169039 |
|  | 07Herpestidae | *Herpestes_pulverulentus* | F | AMNH169040 |
|  | 07Herpestidae | *Herpestes_pulverulentus* | M | cas28740 |
|  | 07Herpestidae | *Herpestes_pulverulentus* | M | lacm30761 |
|  | 07Herpestidae | *Helogale_hirtula* | F | AMNH179103 |
|  | 07Herpestidae | *Helogale_hirtula* | F | AMNH179105 |
|  | 07Herpestidae | *Helogale_hirtula* | M | AMNH179099 |
|  | 07Herpestidae | *Helogale_hirtula* | M | AMNH179100 |
|  | 07Herpestidae | *Helogale_parvula* | F | LACM45747 |
|  | 07Herpestidae | *Helogale_parvula* | F | lacm56754 |
|  | 07Herpestidae | *Helogale_parvula* | M | UWBM35890 |
|  | 07Herpestidae | *Helogale_parvula* | M | UWBM36467 |
|  | 07Herpestidae | *Herpestes_auropunctatus* | F | AMNH163177 |
|  | 07Herpestidae | *Herpestes_auropunctatus* | F | AMNH27600 |
|  | 07Herpestidae | *Herpestes_auropunctatus* | M | FMNH57587 |
|  | 07Herpestidae | *Herpestes_auropunctatus* | M | FMNH75848 |
|  | 07Herpestidae | *Herpestes_brachyurus* | F | FMNH88604 |
|  | 07Herpestidae | *Herpestes_brachyurus* | M | FMNH43343 |
|  | 07Herpestidae | *Herpestes_brachyurus* | M | FMNH88603 |
|  | 07Herpestidae | *Herpestes_edwardsii* | F | AMNH70232 |
|  | 07Herpestidae | *Herpestes_edwardsii* | F | FMNH83097 |
|  | 07Herpestidae | *Herpestes_edwardsii* | F | FMNH97850 |
|  | 07Herpestidae | *Herpestes_edwardsii* | M | FMNH97848 |
|  | 07Herpestidae | *Herpestes_edwardsii* | M | FMNH97852 |
|  | 07Herpestidae | *Herpestes_fuscus* | F | FMNH95035 |
|  | 07Herpestidae | *Herpestes_fuscus* | F | FMNH99527 |
|  | 07Herpestidae | *Herpestes_fuscus* | M | FMNH92886 |
|  | 07Herpestidae | *Herpestes_fuscus* | M | FMNH96327 |
|  | 07Herpestidae | *Herpestes_ichneumon* | F | AMNH118874 |
|  | 07Herpestidae | *Herpestes_ichneumon* | F | AMNH51588 |
|  | 07Herpestidae | *Herpestes_ichneumon* | M | AMNH82779 |
|  | 07Herpestidae | *Herpestes_ichneumon* | M | lacm30317 |
|  | 07Herpestidae | *Herpestes_javanicus* | F | AMNH101656 |
|  | 07Herpestidae | *Herpestes_javanicus* | F | AMNH101658 |
|  | 07Herpestidae | *Herpestes_javanicus* | M | FMNH106498 |
|  | 07Herpestidae | *Herpestes_javanicus* | M | FMNH39349 |
|  | 07Herpestidae | *Herpestes_naso* | M | FMNH25308 |
|  | 07Herpestidae | *Herpestes_naso* | M | FMNH43738 |
|  | 07Herpestidae | *Herpestes_sanguineus* | F | LACM35665 |
|  | 07Herpestidae | *Herpestes_sanguineus* | F | LACM56709 |
|  | 07Herpestidae | *Herpestes_sanguineus* | M | LACM36687 |
|  | 07Herpestidae | *Herpestes_sanguineus* | M | LACM56708 |
|  | 07Herpestidae | *Herpestes_semitorquatus* | F | FMNH88605 |
|  | 07Herpestidae | *Herpestes_semitorquatus* | M | FMNH74274 |
|  | 07Herpestidae | *Herpestes_smithii* | F | AMNH163179 |
|  | 07Herpestidae | *Herpestes_smithii* | F | AMNH171168 |
|  | 07Herpestidae | *Herpestes_urva* | F | AMNH112747 |
|  | 07Herpestidae | *Herpestes_urva* | F | AMNH163605 |
|  | 07Herpestidae | *Herpestes_urva* | M | AMNH60151 |
|  | 07Herpestidae | *Herpestes_urva* | M | FMNH39351 |
|  | 07Herpestidae | *Herpestes_vitticollis* | M | AMNH163180 |
|  | 07Herpestidae | *Herpestes_vitticollis* | M | AMNH240922 |
|  | 07Herpestidae | *Ichneumia_albicauda* | F | AMNH187765 |
|  | 07Herpestidae | *Ichneumia_albicauda* | F | AMNH33316 |
|  | 07Herpestidae | *Ichneumia_albicauda* | M | FMNH98949 |
|  | 07Herpestidae | *Ichneumia_albicauda* | M | lacm40138 |
|  | 07Herpestidae | *Mungos_mungo* | F | AMNH118859 |
|  | 07Herpestidae | *Mungos_mungo* | F | AMNH51127 |
|  | 07Herpestidae | *Mungos_mungo* | M | LACM53757 |
|  | 07Herpestidae | *Mungos_mungo* | M | LACM56719 |
|  | 07Herpestidae | *Paracynictis_selousi* | F | MVZ88710 |
|  | 07Herpestidae | *Suricata_suricatta* | F | AMNH161604 |
|  | 07Herpestidae | *Suricata_suricatta* | F | LACM41576 |
|  | 07Herpestidae | *Suricata_suricatta* | M | LACM56689 |
|  | 07Herpestidae | *Suricata_suricatta* | M | usnm384037 |

Table S2. Comparisons of the best-fitting evolutionary models in cranial shape and mandibular shape. Small sample–corrected Akaike weights (AICcW) were calculated for each of the 250 replications to account for uncertainty in phylogenetic topology and the ancestral character states. Rows in boldface type represent the best-fit model as indicated by ΔAICc < 2. ΔAICc=the mean of AICc minus the minimum AICc between models.

|  | Model | AICc | ∆AICc | AICcW |
| --- | --- | --- | --- | --- |
| A. Cranial shape | |  |  |  |
|  | **mvBM1** | **-702.42** | **0.00** | **0.66** |
|  | mvOU1 | -695.87 | 6.55 | 0.02 |
|  | **mvOUM_felid-fossa_** | **-700.95** | **1.47** | **0.32** |
|  | **mvBM1** | **-702.42** | **0.20** | **0.47** |
|  | mvOU1 | -695.87 | 6.75 | 0.02 |
|  | **mvOUM_viverrid-euplerines_** | **-702.62** | **0.00** | **0.52** |
|  | **mvBM1** | **-702.42** | **0.00** | **0.96** |
|  | mvOU1 | -695.87 | 6.55 | 0.04 |
|  | mvOUM_viverrid-fossa_ | -638.54 | 63.88 | 0.00 |
|  | mvBM1 | -702.42 | 7.97 | 0.02 |
|  | mvOU1 | -695.87 | 14.52 | 0.00 |
|  | **mvOUM_herpestid-galidiines_** | **-710.39** | **0.00** | **0.98** |
| B. Mandibular shape | |  |  |  |
|  | mvBM1 | -1183.91 | 126.97 | 0.00 |
|  | **mvOU1** | **-1310.88** | **0.00** | **0.98** |
|  | mvOUM_felid-fossa_ | -1302.55 | 8.33 | 0.02 |
|  | mvBM1 | -1183.91 | 126.97 | 0.00 |
|  | **mvOU1** | **-1310.88** | **0.00** | **0.99** |
|  | mvOUM_viverrid-euplerines_ | -1301.42 | 9.46 | 0.01 |
|  | **mvBM1** | **-1026.0378** | **0** | **1** |
|  | mvOU1 | -949.99 | 76.04 | 0.00 |
|  | mvOUM_viverrid-fossa_ | -949.01 | 77.03 | 0.00 |
|  | mvBM1 | -1183.91 | 126.97 | 0.00 |
|  | **mvOU1** | **-1310.88** | **0.00** | **0.81** |
|  | mvOUM_herpestid-galidiines_ | -1308.03 | 2.85 | 0.19 |
